# BiomiX 3.0: A user-friendly platform for democratized multi-omics integration with graph-based learning

**DOI:** 10.64898/2026.09.22.753185

**Authors:** Iñigo Clemente-Larramendi, Jessica Gliozzo, Lourdes Velo-Suárez, Ozvan Bocher, Nathan Foulquier, María Hernández-Valladares, Marc-André Legault, Divi Cornec, Christophe Jamin, Alberto Gil-de-la-Fuente, Álvaro Fernández-Ochoa, Cristian Iperi

## Abstract

**Background:** Multi-omics integration has emerged as a powerful strategy to decode the molecular complexity of biological systems. However, the diversity of available methods, each designed with distinct assumptions, objectives, and computational requirements, makes method selection, usage and interpretation challenging for non-expert users. Here we present BiomiX 3.0, an updated version of the BiomiX platform that extends its integration capabilities with four additional methods: Similarity Network Fusion (SNF), NEighborhood-based Multi-Omics clustering (NEMO), Data Integration Analysis for Biomarker discovery using Latent variable approaches for Omics studies (DIABLO), and PRAMIGO (Phenotyping netwoRk Application for Multi-omics InteGratiOn), a novel supervised heterogeneous graph transformer (HGT) introduced in this work.

**Results:** We benchmarked all five methods, MOFA, DIABLO, SNF, NEMO, and PRAMIGO, on two independent multi-omics datasets derived from a Chronic Lymphocytic Leukemia (CLL) cohort comparing IGHV-mutated and unmutated patients, and a pulmonary tuberculosis (PTB) cohort versus healthy controls. Supervised methods (DIABLO, PRAMIGO) consistently achieved higher condition-specific discrimination as measured by the Adjusted Rand Index (ARI) and the Adjusted Mutual Information (AMI). In contrast, unsupervised methods (SNF, NEMO) revealed alternative patient stratifications driven by independent sources of biological variance while MOFA performed in a semi-supervised way occupies an intermediate position, capturing latent factors that explain both disease-associated and orthogonal sources of variance. Systematic gene-centric analysis of the top-ranked features prioritized by each method was supported by manual biological annotation of shared and method-specific signals. Across cohorts, we annotated 154 shared features (116 genes, 38 metabolites) and 60 method-unique features per cohort, demonstrating that no single integration strategy captures the full landscape of biologically relevant signals. In the CLL cohort, shared features spanned B-cell receptor biology, innate immune signaling, RAS/MAPK activation, and epigenetic regulation, while method-unique features revealed supervised-method-specific insights into vesicle trafficking (DIABLO), immune checkpoints (MOFA), and ncRNA regulation (PRAMIGO). In the PTB cohort, a convergent interferon/innate immune signature dominated shared features across all methods. Still, method-unique analysis uncovered DIABLO-specific acylcarnitine metabolic reprogramming, MOFA-specific restoration of lysosomal trafficking, and PRAMIGO-specific γδ T-cell and immunoglobulin repertoire diversity. Across both cohorts, SNF and NEMO proved useful for detecting biological and technical sources of variation that were orthogonal to the primary condition of interest. Ultimately, PRAMIGO uniquely enables the construction of heterogeneous graphs modeling cross-modal molecular interactions, uncovering epigenetic co-regulation programs in CLL and multi-omics inflammatory modules in PTB that are difficult to identify using conventional integration approaches.

**Conclusions:** BiomiX 3.0 provides a graphical user interface (GUI) multi-method integration environment that democratizes access to state-of-the-art multi-omics analysis. By combining both unsupervised and supervised integration strategies within a unified platform and introducing graph-based learning through PRAMIGO, BiomiX 3.0 enables researchers across disciplines with complementary tools to interrogate the biological sources of variation in their data, without requiring bioinformatics expertise.

## Introduction

The advent of high-throughput technologies has enabled the systematic characterization of biological systems across multiple molecular layers. This paradigm shift is exemplified by multi-omics biobanks like Religious Order Study and Memory and Aging Project (ROSMAP)^1^ in Alzheimer or The Cancer Genome Atlas Program (TCGA)^2^ in cancer. While such efforts highlight the importance of integrating heterogeneous data modalities, developing effective and accessible methods for multi-omics integration comes with several computational challenges.

Current strategies for multi-omics integration are commonly categorized as early, intermediate, and late approaches. Early integration concatenates all features into a unified matrix, t tipically requiring a separate step of missing data imputation and often neglecting modality-specific data distributions. Intermediate integration methods address this limitation by modeling each omic layer according to its statistical properties before joint analysis^3^, making them among the most widely adopted frameworks. Late integration analyzes each omics layer independently before combining findings across modalities. Orthogonally, integration methods can be classified as unsupervised, optimized for discovery of intrinsic data structure and patient stratification, or supervised, targeting predefined predictive tasks such as biomarker discovery or outcome prediction^4–6^.

Correlation-based methods such as canonical correlation analysis (CCA)^7^ maximize shared variation across omics layers, with supervised extensions such as DIABLO^8^ further incorporating prediction of a response variable. Matrix factorization approaches decompose multi-omics data into latent factors that capture shared and modality-specific structure, including Multi-Omics Factor Analysis (MOFA)^9,10^ which adds a probabilistic Bayesian framework. Sample-similarity methods instead construct modality-specific graphs in which nodes represent samples and weighted edges encode pairwise similarity. Similarity Network Fusion (SNF)^11^ integrates these graphs through iterative cross-network diffusion to produce a fused sample-similarity representation. NEMO^12^ uses neighborhood-normalized similarities for data fusion and can accommodate samples lacking entire omics modalities without imputation. Deep learning approaches, particularly Graph Neural Networks (GNNs), have recently emerged as flexible frameworks capable of capturing nonlinear cross-modal relationships^13,14^. However, existing GNN-based methods operate primarily on homogeneous graphs with a single node type; heterogeneous graphs, which jointly represent multiple biological entity types and cross-modal relationships, remain underexplored in multi-omics integration, despite their potential to improve interpretability of multi-omics interactions^15^.

Despite this methodological diversity, each available tool was originally designed for a specific task, disease subtyping, biomarker prediction, or factor interpretation. It additionally requires computational expertise in bioinformatics that limits accessibility for biologists and clinicians. Only a handful of platforms, most notably MixOmics^16^, have attempted to democratize these methods, but programming proficiency in R remains required. To the best of our knowledge, none provides a unified environment that integrates single-omics analysis and multiple integration strategies within a single graphical user interface (GUI).

To address this need, we previously developed BiomiX^17^, a GUI desktop platform that enables single-omics analysis of transcriptomics, metabolomics, and methylomics data and their integration through MOFA. Here, we present BiomiX 3.0, which substantially expands the platform’s integration capabilities by incorporating SNF, NEMO, DIABLO and PRAMIGO, a novel supervised heterogeneous graph transformer (HGT) framework introduced in this study to uncover biologically meaningful cross-modal multi-omics programs and regulatory interactions. We benchmarked all five methods across two independent multi-omics datasets and show that each method captures distinct, partially overlapping biological signals, demonstrating that method choice has meaningful consequences for biological discovery. PRAMIGO, by modeling cross-modal molecular interactions through heterogeneous graphs, provides a unique analytical perspective that complements and extends conventional integration approaches. BiomiX 3.0 makes all these tools accessible in a single, user-friendly environment that requires minimal bioinformatics expertise. To further support accessibility, step-by-step tutorials for running BiomiX 3.0 are available on the project website [https://ixi-97.github.io/].

## Methods

### Datasets

#### Chronic Lymphocytic Leukemia (CLL) cohort

The CLL dataset was downloaded from the PACE repository (http://pace.embl.de/)^18^ and comprises whole-blood transcriptomic (RNA-seq) and methylomic (Illumina 450K array) profiles from patients with IGHV-mutated (M-CLL) and IGHV-unmutated (U-CLL) chronic lymphocytic leukemia. Transcriptomic data were normalized using a variance-stabilizing transformation (VST), while methylomic data were represented as beta values. For each omics modality, the 5,000 features with the highest variance were retained, and the resulting transcriptomic and methylomic matrices were used as input for the BiomiX 3.0 integrations, as implemented in BiomiX.

#### Pulmonary Tuberculosis (PTB) cohort

The PTB dataset (ENA accession PRJNA971365)^19^ includes whole-blood transcriptomics and plasma metabolomics profiles from patients with pulmonary tuberculosis (PTB) and healthy controls (HC). Raw FastQ files were processed as described in the original BiomiX publication^17^, using STAR v2.7.11 for alignment, HTSeq-count for gene quantification, and the BiomiX metabolomics pipeline for peak annotation and statistical analysis. Transcriptomic and metabolomic data were normalized using VST and log transformation , respectively. For transcriptomics data, the 5000 features with highest variance were filtered.

### Reproducibility and platform stability

#### Docker system and Nextflow pipeline

BiomiX 3.0 adopts a multi-container architecture in which each single-omics module and multi-omics integration algorithm runs within a dedicated Docker container, minimising cross-dependency conflicts and ensuring version-locked reproducibility across platforms (Ubuntu 24.04, R 4.5.2 Python 3.8). Installation is handled by an automated setup script. The GUI has been migrated from PyQt5 to R Shiny to ensure compatibility with containerised execution. A Nextflow pipeline [https://github.com/BiomiX-consortium/biomix_nextflow_workflow] enables full automation of the BiomiX 3.0 workflow from a single JSON configuration file, supporting all integration methods and making the analysis accessible to users operating on HPC or cloud infrastructures without GUI interaction.

### Multi-omics integration methods

#### MOFA

Multi-Omics Factor Analysis (MOFA)^10^ is an unsupervised probabilistic framework that decomposes multi-omics datasets into a set of latent factors capturing shared and modality-specific sources of variance. In BiomiX 3.0, MOFA is implemented with automated factor selection: models are iteratively trained with increasing numbers of factors, stopping when at least 3 consecutive models show that the final factor explains less than 1% of the total variance. The three best-performing models, assessed by the Mann-Whitney test with FDR correction, are retained. This way, MOFA was implemented in a semi-suipervised way meaning that the best-performing models are the ones leading to better groups discrimination. Discriminant factors are interpreted through correlation with clinical data, pathway enrichment via EnrichR, and PubMed bibliography search, as described in BiomiX.

#### DIABLO

DIABLO (Data Integration Analysis for Biomarker discovery using Latent variable approaches for Omics studies)^8^ is a supervised integration method from the MixOmics^16^ framework that extends sparse canonical correlation analysis to multiple omics layers. It identifies correlated latent components across modalities while optimizing discrimination between predefined groups. In BiomiX 3.0, DIABLO is implemented with automated component number optimization: models are iteratively trained with increasing numbers of components, and the optimal number is selected based on the count of discriminant components and their statistical significance, assessed by the Mann-Whitney test with FDR correction, mirroring the factor selection strategy applied to MOFA. L1 regularization is applied for sparse feature selection. The method requires complete samples across all omics layers; missing samples are excluded before analysis.

#### SNF

Similarity Network Fusion (SNF)^11^ is an unsupervised network-based integration method that constructs a patient similarity network for each omic layer and iteratively fuses them into a single integrated network through a message-passing algorithm. The fused network captures complementary information across omics by allowing each modality to inform the structure of the others. Spectral clustering is applied to the fused network to identify patient subgroups. In BiomiX 3.0, SNF is implemented using the SNFtool R package^11^, with the number of clusters selected automatically via the eigen-gap approach.

#### NEMO

NEMO (NEighborhood-based Multi-Omics clustering)^12^ is a network-based unsupervised method that handles datasets with partially overlapping samples across omics layers without requiring imputation. For each omic separately, NEMO calculates pairwise sample similarities using a locally scaled radial basis function kernel. These similarities are then normalized relative to each sample’s neighbourhood to obtain omic-specific relative-similarity matrices. The matrices are integrated by averaging, for each pair of samples, only across the omics measured in both samples, thereby avoiding imputation of missing modalities. Z-score was applied before integration. The resulting multi-omics affinity matrix is then used for spectral clustering and the optimal number of clusters is selected via a modified eigengap method. In BiomiX 3.0, NEMO is implemented using the NEMO R package.

#### PRAMIGO

PRAMIGO (Phenotyping netwoRk Application for Multi-omics InteGratiOn) is a novel supervised Heterogeneous Graph Transformer (HGT)^20,21^ introduced in this study. PRAMIGO combines autoencoder-based representation learning with heterogeneous graph transformers to model cross-modal biological interactions. First, each omic layer and the sample matrix are independently compressed into a shared latent space using three autoencoders. These generate low-dimensional embeddings for omic features and samples, enabling direct comparison between heterogeneous entities. The autoencoders are optimized by minimizing reconstruction error using mean squared error (MSE):

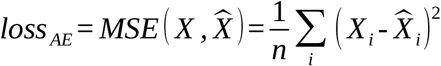

Where *n* is the number of samples. The latent embeddings are then used to construct a heterogeneous graph *G* =(*V* , *E* , *A* , *R*), where *V* denotes the set of nodes, *E* the set of edges, *A* the adjacency structure, and *R* the set of relation types. Nodes represent samples and omic features, whereas edges encode within-omic, cross-omic, and sample-omic relationships. Within-omic edges are inferred using Pearson correlation between feature embeddings, whereas cross-modal relationships are established using K-nearest neighbor (KNN) graphs based on Euclidean distance in latent space. The resulting graph is processed using a Heterogeneous Graph Transformer (HGT), which performs relation-aware attention-based message passing across node and edge types. For each target node *v*_*t*_, attention weights are computed over neighboring source nodes *v*_*s*_ using type-specific query, key and value projections:

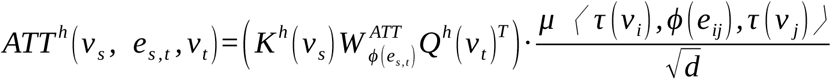

In this formulation, *v*_*s*_and *v*_*t*_denote the source and target nodes, respectively, while *e*_*s,t*_represents the edge connecting them. *K*^*h*^ (*v*_*s*_) and *Q*^*h*^ (*v*_*t*_) are the type-specific key and query representations of the source and target nodes for attention head *h*, respectively. 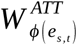 encodes an edge-type-specific transformation based on the relation *ϕ* (*e*_*s* , *t*_), while 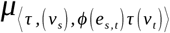 is a learnable parameter associated with the complete meta-relation defined by the source node type, edge type, and target node type. Finally, *d*denotes the dimensionality of the node representations and provides the scaling factor used in the attention computation. Messages from neighboring nodes are aggregated to update node embeddings across HGT layers, enabling the model to learn biologically relevant cross-modal interactions.

PRAMIGO is trained using a supervised representation-learning objective that combines a cross-entropy classification loss, *L*_*CE*_, with a cosine embedding loss, *L*_cos_

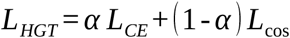

The cross-entropy term optimizes the prediction of sample conditions or classes, ensuring that the learned representations retain discriminative information relevant to the supervised task. The cosine embedding loss further structures the latent space by encouraging samples belonging to the same condition to exhibit similar embeddings while increasing the separation between samples from different conditions. The relative contribution of the two objectives is controlled by the weighting parameter *α*. This architecture enables PRAMIGO to identify condition-specific multi-omics network programs and biologically interpretable feature interactions. Full architectural details are provided in the Results section and in Supplementary Methods.

### State-of-the-art graph neural network benchmarking

We benchmarked PRAMIGO against the state-of-the-art multi-omics graph neural network methods MOGONET^13^ and HeteroGATomics^15^. Following the evaluation framework of HeteroGATomics^15^, we downloaded three publicly available cancer datasets from The Cancer Genome Atlas: bladder urothelial carcinoma (BLCA), lower-grade glioma (LGG) and renal cell carcinoma (RCC), each integrating DNA methylation, mRNA and miRNA profiles. The BLCA and LGG datasets were formulated as binary classification tasks, distinguishing high-grade from low-grade tumors in BLCA and Grade II from Grade III tumors in LGG. In contrast, RCC was formulated as a three-class classification task, distinguishing kidney chromophobe carcinoma (KICH), kidney renal clear cell carcinoma (KIRC), and kidney renal papillary cell carcinoma (KIRP).

Classification performance was evaluated using stratified 10-fold cross-validation. For the binary classification tasks (BLCA and LGG), performance was assessed using the Area Under the Receiver Operating characteristic Curve (AUROC), accuracy, Positive Predictive Value (PPV; precision), Negative Predictive Value (NPV), sensitivity (recall), and specificity. For the multiclass RCC task, performance was evaluated using accuracy, precision, recall, and macro-, micro-, and weighted-F1 scores. When AUROC and AUPRC were computed for the multiclass RCC task, multiclass performance was calculated using a one-versus-rest strategy and aggregated across classes using the specified averaging scheme.

### Benchmarking metrics

To evaluate the performance of all five integration methods comparably and objectively, we applied a standardized set of metrics covering three complementary analytical dimensions.

The agreement between method-derived cluster assignments and known condition labels (called condition conservation) was assessed using the Adjusted Rand Index (ARI) and adjusted mutual information (AMI), which measure the agreement between method-derived cluster assignments and known condition labels. These metrics are corrected for chance and range from 0 (random agreement) to 1 (perfect agreement). For supervised methods (DIABLO, PRAMIGO), cluster assignments were derived from latent-space representations using k-means clustering (k = number of conditions).

Intrinsic cluster quality was evaluated using the Average Silhouette Width (ASW) and the Calinski-Harabasz (CH) index, which measure within-cluster cohesion and between-cluster separation independently of known labels. These metrics are particularly informative for assessing whether unsupervised methods identify novel, internally consistent patient stratifications.

On top of standard metrics, condition separation in latent space was quantified as the ratio of the mean inter-condition to intra-condition Euclidean distances, providing an intuitive measure of how clearly the two conditions are resolved in the learned representation.

Space visualization of PRAMIGO, DIABLO, and MOFA was performed using UMAP, tSNE and PHATE as these methods compute a latent space, while NEMO, and SNF using Isomap, MDS and Spectral as these methods calculate a similarity matrix.

Relative contribution analysis was performed using the different sources of weighting information in each method: DIABLO’s global variance explained by omic; MOFA’s variance explained by each omic in each factor; SNF’s and NEMO’s features importance score of the top 100 omic features from each omic through Mutual Information (MI) analysis (similarly to SNFtool rankFeaturesByNMI function); HGT’s attention score for all the omic features. Full calculation details are provided in Supplementary Methods.

Feature overlap across methods was assessed using the Jaccard Index computed over the top 20 features (by absolute loading weight or attention score and NMI score) identified by each method, separately for each omic layer. This number of feature was selected to facilitate interpretability and it is set as default hyperparameterin BiomiX 3.0.

NEMO, SNF and PRAMIGO were applied only on samples with both omics available. Full hyper-parametrization of the methods is described in Supplementary Methods.

### Feature-level biological signal analysis

To characterize the biological content captured by each method independently of formal pathway enrichment statistics, we adopted a two-step feature-centric approach. In the first step, feature prioritization and overlap analysis, the top contributing genes and metabolites were extracted from each method based on absolute loading weights (MOFA, DIABLO) or feature importance scores (SNF, NEMO, PRAMIGO). For MOFA, only factors significantly discriminating between conditions (Mann-Whitney FDR < 0.05) were considered; for DIABLO, features were extracted from all significant latent components. Methylomics CpG sites were mapped to their associated genes where HGNC annotation was available. Feature overlaps between methods were computed separately for the top features from each omic layer, and features detected by two or more methods were classified as ‘shared’. In contrast, those detected by exactly one method were classified as ‘method-unique’. This analysis was implemented in R, with results were stored as CSV files (Supplementary Tables S1-S2).

In the second step, biological annotation, shared and method-unique features were manually annotated using a disease-context-aware curation strategy. For each gene, the primary biological process was determined based on curated databases (Gene Ontology, UniProt, HGNC) and disease-specific literature for CLL and PTB. For metabolites, compound identities were resolved from HMDB identifiers using the Human Metabolome Database reference file, and each metabolite was assigned to a biological class (lipid metabolism, nucleotide metabolism, amino acid/tryptophan catabolism, or xenobiotic) based on its known biochemical role. Features were grouped into biologically coherent categories, and each category was assigned a distinct color for visualization.

Specifically, for the CLL cohort, features were manually annotated into nine biological categories: B-cell and IG, Epigenetic Regulation, ECM and Cytoskeleton, Membrane and Receptor, Lipid and Metabolism, Ubiquitin and Proteostasis, and Other. For the PTB cohort, features spanning both RNA and metabolomics were organised into ten categories: Interferon and Innate Immunity, NK and Cytotoxic T Cells, T-cell and Adaptive Immunity, Erythroid and O_2_ Transport, Granuloma and Monocyte Trafficking, Lipid and Metabolic Reprogramming, Nucleotide Metabolism, Tryptophan and AA Metabolism, Xenobiotic and Drug, and Other. The Lipid and Metabolic Reprogramming category in the PTB cohort was the only category to contain features from both omic layers simultaneously, encompassing phosphatidylcholine-PUFA species, lysophosphatidylcholines, and lipid-related genes, making the RNA-metabolome convergence on lipid dysregulation visually explicit in Fig. 3 layout. A complete list of biological categories and their constituent features for both cohorts is provided in Supplementary Table S3 (CLL) and Supplementary Table S4 (PTB).

### PRAMIGO multi-omics programs analysis

Epigenetic programs identified in the CLL dataset were analyzed using a correlation matrix of epigenetic feature embeddings, revealing correlated hubs. Multi-omics programs identified in the PTB dataset were performed by isolating the attention network from the top 100 transcriptomic and metabolomic features. On that multi-omics weighted attention network, we performed community detection using Leiden clustering^32^ with resolution 1. We kept the modules with more than 2 elements.

## Results

### BiomiX 3.0: an expanded platform for multi-omics integration

The updated version of BiomiX maintains full compatibility with the single-omics analysis pipelines described in BiomiX v1^17^, including transcriptomics (DESeq2/Limma), metabolomics (CEU Mass Mediator, TidyMass), and methylomics (ChAMP), and extends the integration module with four additional methods: SNF, NEMO, DIABLO, and PRAMIGO (**Fig. 1A**). All methods are accessible through the same graphical user interface (GUI) accepting standard matrix inputs and producing structured output reports with figures and interpretable results files. The platform continues to support automated MOFA factor number optimization, clinical data correlation, pathway analysis via EnrichR, and PubMed bibliography search for factor annotation.

**Fig. 1:**
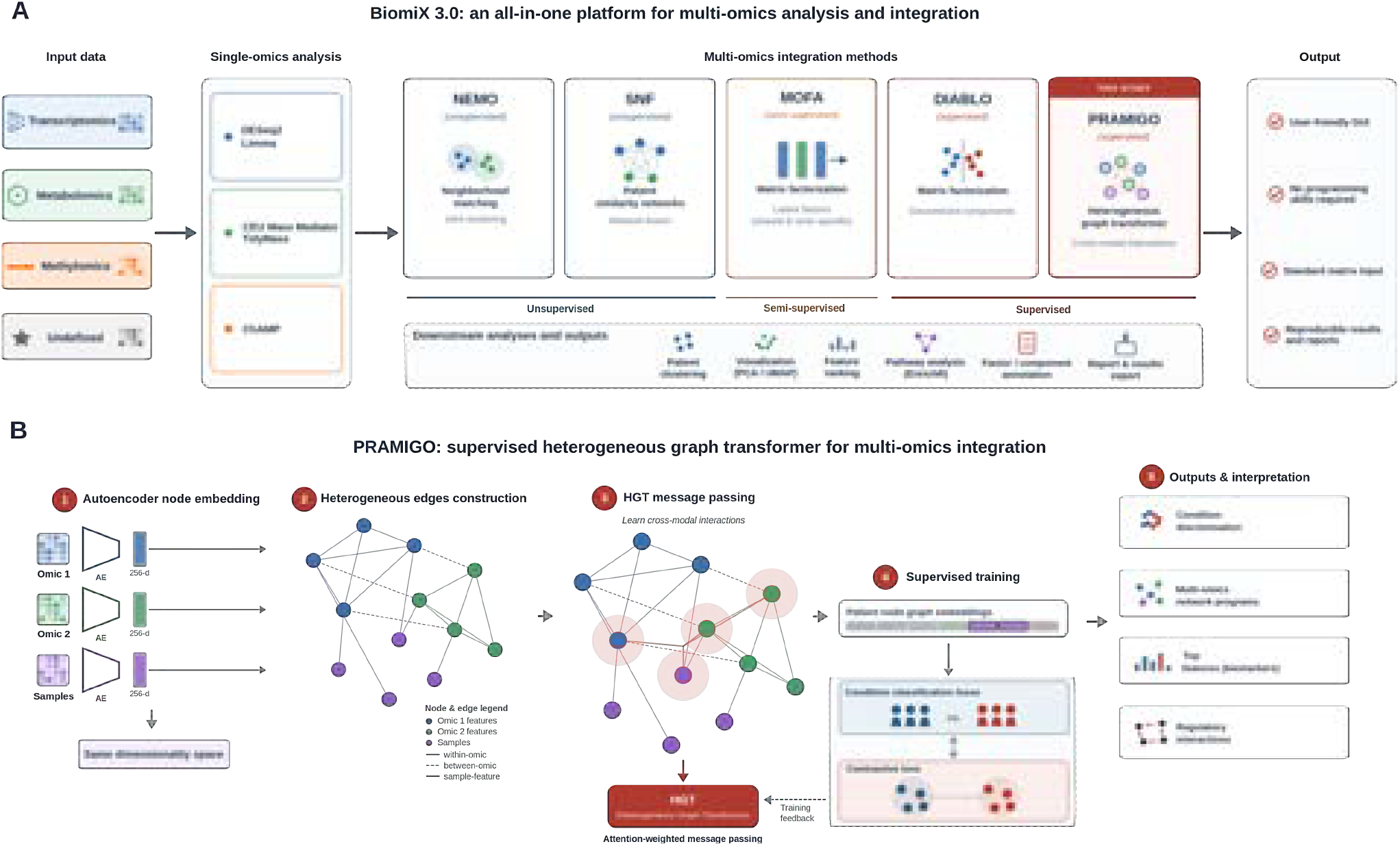
BiomiX 2.0 platform overview. (A) Schematic of the BiomiX 3.0 pipeline showing single-omics analysis modules (transcriptomics, metabolomics, methylomics) and the five available integration methods. (B) PRAMIGO heterogeneous graph architecture showing the three node types (samples, omic-1 features, omic-2 features), three edge classes (within-omic, between-omic, sample-feature), autoencoder embedding, HGT message passing, and contrastive loss training.

The five integration methods span two orthogonal axes of methodological diversity. On the supervision axis, MOFA, SNF, and NEMO are unsupervised and do not require prior knowledge of group labels. At the same time, DIABLO and PRAMIGO are supervised and optimize integration toward condition discrimination. On the modeling axis, MOFA and DIABLO use matrix factorization to extract latent components, whereas SNF and NEMO operate on patient-similarity networks. PRAMIGO learns directly from a heterogeneous graph that jointly represents samples and molecular features (**Fig. 1B**). This diversity of analytical frameworks is a deliberate design choice: rather than selecting a single best-performing method, BiomiX 3.0 provides complementary perspectives on multi-omics data, allowing users to interrogate different aspects of biological variation.

PRAMIGO is a supervised graph neural network framework designed to identify biologically meaningful multi-omics patterns that distinguish two conditions. The method integrates multiple omics datasets by constructing a heterogeneous graph in which nodes represent both samples and molecular features, while edges capture relationships within and between omics layers. This design allows PRAMIGO to simultaneously model interactions across samples and molecular entities, providing an interpretable representation of cross-modal biological networks.

To build this graph, PRAMIGO first compresses each omic dataset into a shared latent space using autoencoders, enabling direct comparison between different data types. Relationships between nodes are then inferred using correlation- and distance-based metrics to connect biologically related features and samples. The resulting heterogeneous graph is processed using a Heterogeneous Graph Transformer (HGT), which applies attention-based learning to identify the most informative cross-modal interactions associated with the studied condition. Through supervised contrastive training, PRAMIGO learns multi-omics network programs that both discriminate disease states and highlight potential regulatory mechanisms.

### Benchmarking multi-omics integration methods across two independent cohorts

We benchmarked the predictive performance of PRAMIGO against state-of-the-art methods across three independent multi-omics reference datasets (**Fig. 2**). We subsequently compared all five BiomiX 3.0 methods on two independent multi-omics datasets, the CLL cohort (transcriptomics and methylomics) and the PTB cohort (transcriptomics and metabolomics), using a standardized set of metrics covering condition conservation, intrinsic cluster quality, latent space separation, and feature prioritization (**Fig. 3**).

**Fig. 2:**
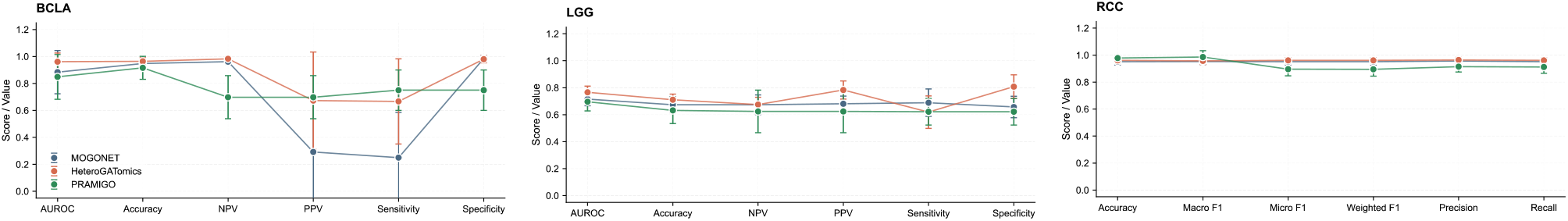
Benchmarking PRAMIGO with state-of-the-art multi-omics integration Graph Neural Networks. AUROC is the Area Under the Receiver Operating Characteristic curve, representing the model’s ability to discriminate between diagnostic classes. Accuracy is the proportion of total instances correctly classified. NPV/PPV are Negative Predictive Value and Positive Predictive Value (Precision), respectively. Sensitivity (Recall) measures the true positive rate, while Specificity measures the true negative rate. F1-Scores (Macro, Micro, Weighted) are the harmonic means of precision and recall calculated at different levels of class aggregation for the RCC dataset.

**Fig. 3:**
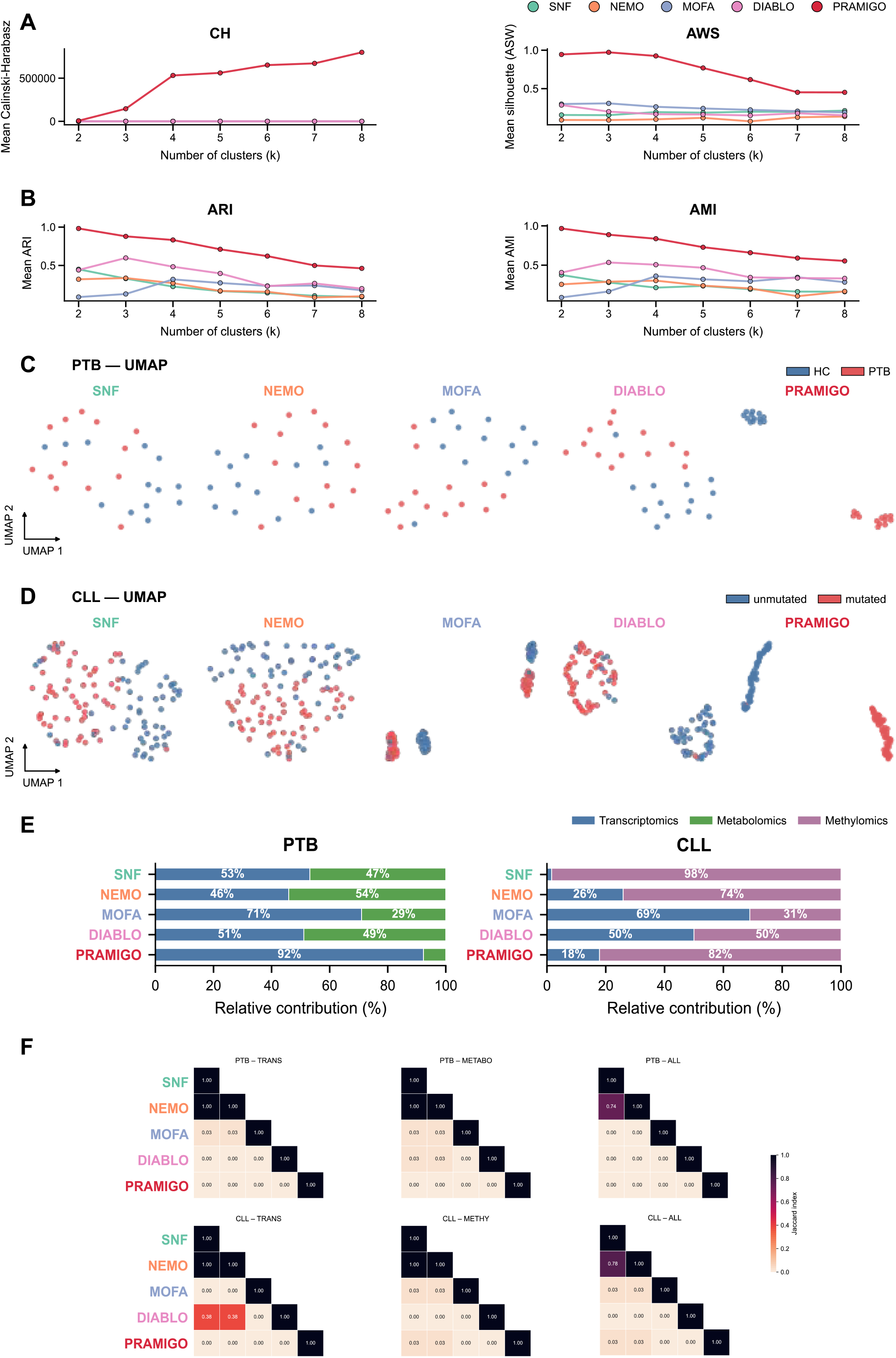
Benchmarking of five multi-omics integration methods on CLL and PTB cohorts. (A) Adjusted Rand Index (ARI) and adjusted mutual information (AMI) for all five methods on both cohorts. (B) Average silhouette width (ASW) and Calinski-Harabasz (CH) index. (C, D) UMAP visualizations of latent embeddings colored by condition label for CLL (C) and PTB (D). (E) Omic relative contribution per method and cohort. (F) Jaccard Index of top-20 feature overlap between all method pairs.

#### Performance relative to existing graph-based methods

We compared PRAMIGO to state-of-the-art graph neural network approaches for multi-omics integration, including MOGONET^13^ and HeteroGATomics^15^ (**Fig. 2**). Benchmarking was performed across three reference cancer datasets (BLCA, LGG, and RCC), each integrating DNA methylation, mRNA, and miRNA profiles, following previously established evaluation settings^20^. Classification performance was assessed using stratified 10-fold cross-validation. PRAMIGO achieves comparable performance over both methods in BLCA, LGG and RCC highlighting better Sensitivity and PPV in BCLA dataset.

#### Condition conservation and latent space separation

Supervised methods consistently achieved higher condition-specific discrimination than unsupervised approaches. In both cohorts, DIABLO and PRAMIGO yielded the highest ARI and AMI values, confirming their ability to preserve known group structure in the learned representation (**Fig. 3A** and **Supplementary Fig. 1A**). PRAMIGO achieved the highest inter/intra-condition distance ratio in both datasets, indicating particularly clear separation of the two conditions in its latent space (**Supplementary Fig. 1C**). MOFA performed intermediately: its discriminant factors successfully separated conditions, but the overall latent space also captured substantial non-condition variance. SNF and NEMO produced mixed result in which known condition labels were not the primary driver of cluster structure (**Fig. 3C, D** and **Supplementary Fig. 2**).

#### Unsupervised methods reveal independent sources of biological variance

Despite lower condition-specific scores, SNF and NEMO revealed alternative patient stratifications, achieving ASW and CH index values comparable to MOFA and higher than those obtained with DIABLO (**Fig. 3B** and **Supplementary Fig. 1B**). In both cohorts, the primary driver of unsupervised clustering was not the condition of interest but an independent source of variance, sex-linked gene expression and age-associated metabolic variation (**Supplementary Fig. 3,4**). This is not a methodological failure: it reflects the unsupervised nature of these approaches, which capture the dominant source of variance in the data irrespective of phenotypic labels. These findings highlight an important practical consideration: the choice of integration method should be guided by the analytical objective. When the goal is to characterize condition-specific molecular mechanisms, supervised methods are preferable. When the goal is to discover novel patient subgroups or identify confounding sources of contributions, unsupervised methods provide irreplaceable complementary information.

#### Feature prioritization and omic contributions

We next examined the omic contribution to the learned representations and the overlap between features prioritized by different methods with the relative contribution analysis (**Fig. 3F**). DIABLO assigned approximately equal weight to both omics layers, whereas MOFA attributed substantially greater variance to transcriptomics. SNF, NEMO, and PRAMIGO dynamically adapted their omic contributions to the dataset. In the CLL cohort, methylomics contributed more prominently across all three methods, consistent with well-established epigenetic differences between M-CLL and U-CLL. Feature overlap, assessed using the Jaccard Index across the top 20 features per method, was low across all method pairs (median Jaccard < 0.15), indicating that the five methods largely prioritize distinct molecular features. Unsupervised methods showed higher inter-method feature overlap than supervised approaches, consistent with their shared focus on dominant sources of variance rather than condition-specific signals.

### Each integration method captures distinct biological signals

The benchmarking metrics described above characterize the statistical properties of each method’s output, but do not directly address the biological content of the identified features. To assess what each method actually captures at the gene and metabolite level, we performed a systematic biological annotation of all top-ranked shared and method-unique features across both cohorts (**Fig. 4** and **Supplementary Tables S1 –S2**). Features detected by two or more methods were classified as ‘shared’; features detected by exactly one method were classified as ‘method-unique’ (top 100 per method per cohort;).

**Fig. 4.**
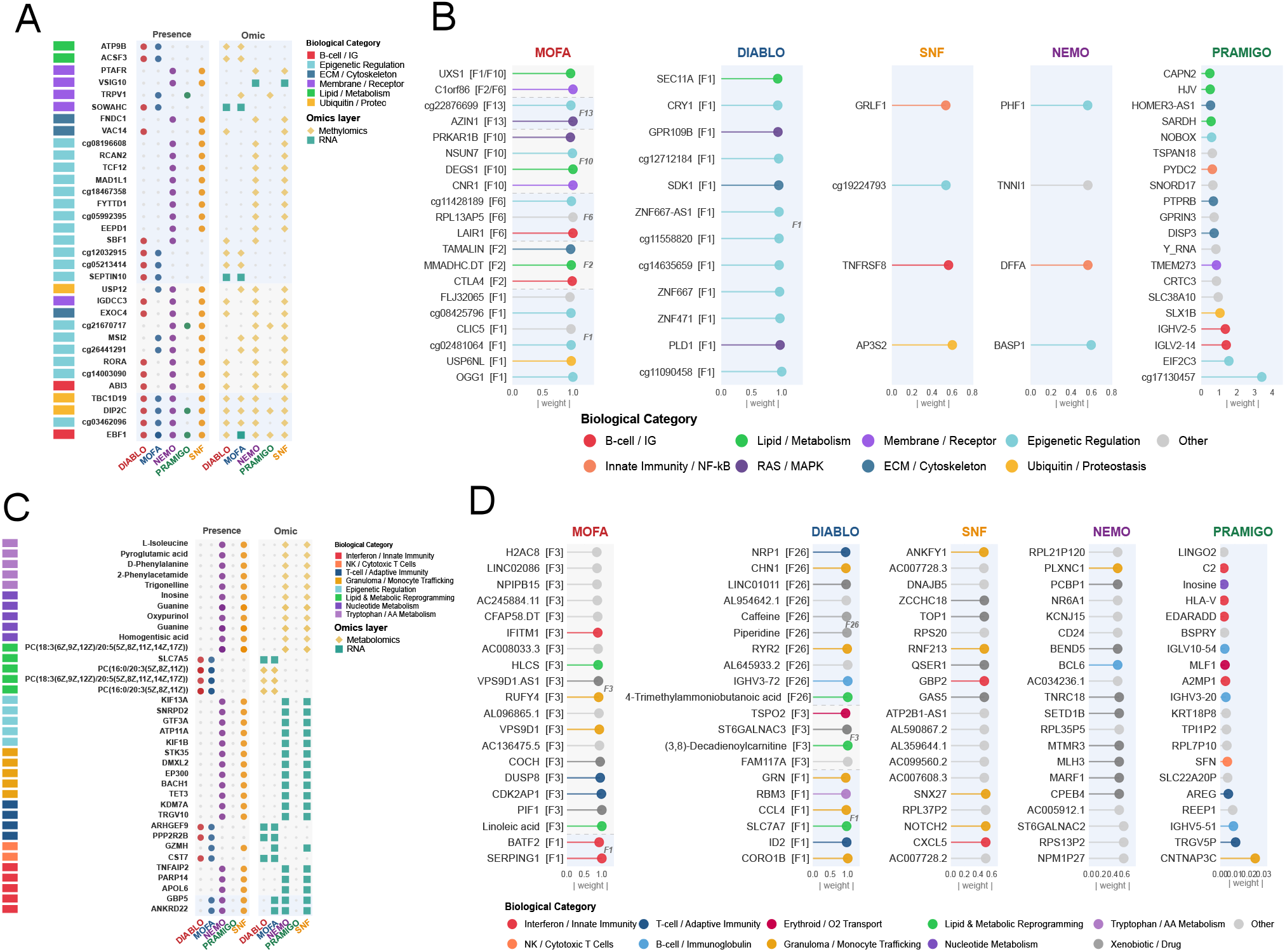
Gene-level biological signal comparison across integration methods. (A-B) CLL cohort. (A) Structured dot plot showing the top contributing genes identified by MOFA (discriminant factors F1, F2, F6, F10, F13), DIABLO (factor 1), SNF, NEMO, and PRAMIGO. Rows represent genes/CpG sites; columns represent methods. Dot size encodes absolute loading weight or importance score. Left sidebar indicates the biological process category (color legend). Right panels show presence/absence matrix and omic layer of origin (RNA: teal square; methylomics: gold diamond). Genes are grouped by overlap tier. (B) Lollipop plot of top-100 method-unique genes per method in CLL, ordered by factor of origin and colored by biological category. (C-D) PTB cohort. (C) Structured dot plot as in (A), with an additional metabolomics omic layer (metabolomics: purple diamond). The ‘Lipid & Metabolic Reprogramming’ category (green) contains both RNA and metabolomics features, making the cross-omic convergence on lipid dysregulation directly visible. (D) Lollipop plot of top-100 method-unique features per method in PTB.

#### CLL cohort: convergent B-cell signals with method-specific epigenetic and signaling depth

In the CLL cohort, 33 features were shared across two or more methods, spanning transcriptomics and methylomics. The shared feature landscape (**Fig. 4A**) was organised into six biological categories, with Epigenetic Regulation constituting the largest group (18/33 features).

Two features were identified by all five methods: *EBF1* (B-cell lineage master transcription factor) and DIP2C (DNA methylation reader). At the four-method level, cg03462096 (*KTN1* locus, IGHV-stratified DMR) and *TBC1D19* (Rab GTPase-activating protein) were co-identified by DIABLO, MOFA, SNF, and NEMO. The Epigenetic Regulation category comprised nine CpG probes, including cg12032915 (*LMBR1* S-shelf, part of the validated nine-CpG IGHV prognostic panel), and nine genes including *RORA, MSI2, TCF12*, and *MAD1L1*, identified across combinations of SNF, NEMO, DIABLO, and MOFA. Additional shared categories included Ubiquitin/Proteostasis (*DIP2C, TBC1D19, USP12*; three to five methods), ECM/Cytoskeleton (*EXOC4, VAC14, FNDC1*), and Membrane/Receptor (*IGDCC3, SOWAHC, TRPV1, VSIG10, PTAFR*).

Analysis of method-unique features (**Fig. 4B** and **Supplementary Table S3**) revealed distinct biological programmes for each method. DIABLO-unique features (n=12) included a KRAB-ZNF repressor cluster (*ZNF471, ZNF667, ZNF667-AS1*), the circadian regulator *CRY1*, the phospholipase *PLD1*, and *SDK1*. MOFA contributed the largest set of unique features (n=120), spanning immune checkpoints (*CTLA4, LAIR1*), DNA repair (*OGG1*), sphingolipid metabolism (DEGS1), and epitranscriptomic regulators (*NSUN7*), consistent with its multi-factor unsupervised decomposition across independent latent dimensions. SNF and NEMO each identified four unique features: *AP3S2, TNFRSF8, cg19224793* and *GRLF1* for SNF; *BASP1, DFFA, TNNI1*, and *PHF1* for NEMO. PRAMIGO-unique features (n=27) were led by immunoglobulin variable region genes (*IGLV2-14*; *IGHV2-5*), the CpG probe cg17130457, and *EIF2C3* (AGO3/RISC complex).

#### PTB cohort: lipid metabolic reprogramming as the convergent shared signal with method-specific immune and metabolic depth

In the PTB cohort (transcriptomics and metabolomics), 134 features were shared across two or more methods and included in ten biological categories. The shared feature landscape (**Fig. 4C**) was dominated by Lipid & Metabolic Reprogramming (52/134 features), comprising an extensive panel of phosphatidylcholines (PC), lysophosphatidylcholines (LysoPC), phosphatidylethanolamines (PE), sphingomyelin, and linoleic acid, co-identified predominantly by SNF and NEMO. Additional shared metabolite categories included Tryptophan/AA Metabolism (10 features: Trigonelline, Ornithine, Pyroglutamic acid, Choline, Betaine, and related metabolites; SNF/NEMO) and Nucleotide Metabolism (Inosine, Guanine, Homogentisic acid, Oxypurinol; SNF/NEMO). A cross-omic lipid signal was also captured by DIABLO and MOFA jointly, with PC(16:0/20:3), PC(18:3/20:5), and *SLC7A5* shared between these two supervised methods.

Among shared transcriptomic features, Interferon/Innate Immunity was represented by *ANKRD22* and *GBP5* (three methods: MOFA, SNF, NEMO) alongside *APOL6, PARP14, TNFAIP2*, and *WDFY1* (SNF/NEMO). T-cell/Adaptive Immunity features included *PPP2R2B* and *ARHGEF9* (DIABLO/MOFA) and *TRGV10* and *KDM7A* (SNF/NEMO). NK/Cytotoxic T Cell markers *CST7* (DIABLO/MOFA) and *GZMH* (MOFA/SNF) were also co-identified. Epigenetic Regulation (12 features, including *KIF1B, KIF13A, ATXN7*, and *AFF1*) and Granuloma/Monocyte Trafficking (5 features: *TET3, BACH1, EP300, DMXL2, STK35*) were shared exclusively between SNF and NEMO.

Analysis of method-unique features (**Fig. 4D** and **Supplementary Table S4**) revealed distinct biological programmes per method. DIABLO identified 81 unique features, including the acylcarnitines (3,8)-Decadienoylcarnitine and 4-Trimethylammoniobutanoic acid, *CORO1B, IGHV3-72*, and xenobiotics Caffeine and Piperidine. MOFA contributed 48 unique features, including *SERPING1, PIF1, CDK2AP1, DUSP8, COCH*, and additional PUFA-containing PC species across independent factors. SNF and NEMO each identified large sets of unique features (75 and 76 respectively), predominantly additional PC/LysoPC lipid species and non-coding RNA loci at low weights. PRAMIGO-unique features (n=30) included Inosine, *CNTNAP3C, TRGV5P, IGHV5-51, AREG*, and *REEP1*.

### PRAMIGO reveals cross-modal molecular programs inaccessible to conventional integration methods

Having established the complementarity of the five methods at the gene level, we next investigated whether the heterogeneous graph architecture of PRAMIGO provides information that cannot be obtained from conventional approaches. Specifically, we asked whether the cross-modal molecular interactions explicitly modeled by PRAMIGO’s attention mechanism reveal biologically coherent programs beyond those identifiable by factorization- or network-based methods. PRAMIGO constructs a heterogeneous graph linking samples and features across two omic modalities, with within- and cross-omic interactions inferred from correlations and nearest-neighbor relationships, and processes this graph using a Heterogeneous Graph Transformer trained end-to-end with supervised contrastive learning to capture condition-discriminant, cross-modal representations.

#### CLL: three epigenetic programs underlying IGHV mutation status

In the CLL cohort, inspection of the highest-attention subgraph revealed that the top-ranked features were predominantly methylomic, with methylomics nodes forming the most densely connected components of the learned graph (**Fig. 5A**). This is consistent with the established importance of epigenetic reprogramming in IGHV-defined CLL subgroups, and suggests that PRAMIGO’s adaptive weighting naturally amplifies the omic layer carrying the strongest condition-discriminant signal.

**Fig. 5:**
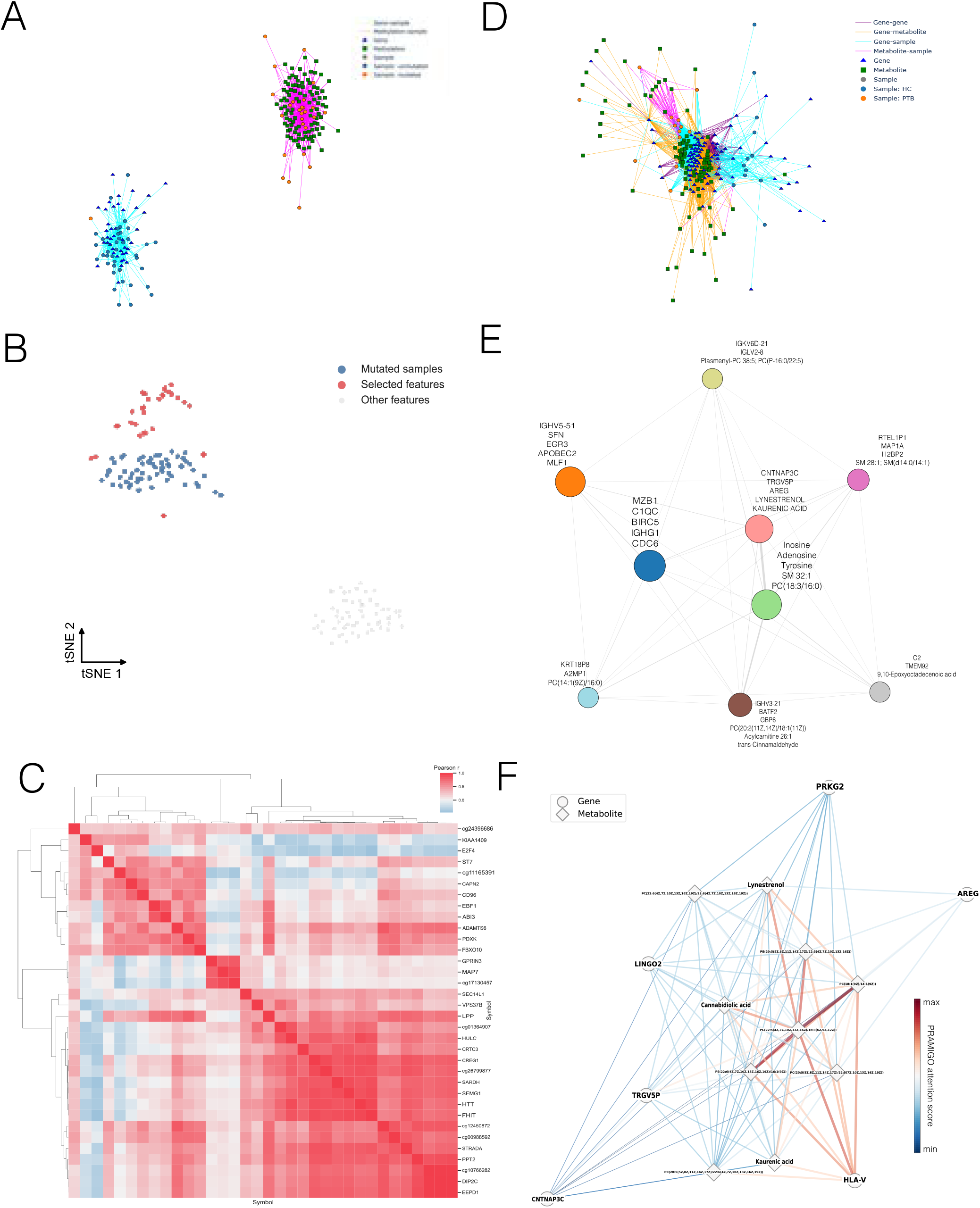
PRAMIGO identifies methylation hubs in mutated CLL and Pulmonary Barrier Disruption & Neuromodulation program in PTB. (A) Attention network of the top 100 genes and top 100 methylomic genes with samples. (B) tSNE of the top 100 methylomic genes embeddings with mutated CLL embeddings. (C) Methylation hubs from a correlation heatmap of the top 34 methylomic gene embeddings. (D) Attention network of the top 100 transcriptomic features and top 100 methylomic features with samples. (E) The Multi-omics Program Network was obtained from Leiden clustering on the isolated top 100 transcriptomic features and top 100 methylomic features attention network. (F) Pulmonary barrier disruption and neuromodulatory program identified by PRAMIGO.

By jointly analyzing methylomic feature embeddings and mutated-sample embeddings, we identified 34 methylomic features with the strongest cross-modal associations. Correlation analysis of the corresponding feature embeddings confirmed clear separation between M-CLL and U-CLL samples (**Fig. 5B** and **Supplementary Fig. 5**). Community detection on the induced subgraph resolved three highly connected hubs that characterize the mutated CLL epigenetic state (**Fig. 5C**).

Program 1 constitutes a B-cell lineage and differentiation hub, featuring transcription factors *EBF1* and *E2F4*, surface markers *CD96* and *ABI3*, and CpG sites (cg11165391, cg24396686) at loci associated with memory B-cell identity. This program likely reflects a more mature, post-germinal center epigenetic state in M-CLL relative to U-CLL. Program 2 represents a prognostic signature centered on *MAP7* and cg17130457, both of which have been reported as specific markers of the stable, indolent M-CLL subgroup^33^. Program 3 encompasses a broad epigenetic maturity program, including structural and signaling genes *LPP, HTT*, and *CRTC3*, the long non-coding RNA *HULC*, and tumor suppressors *FHIT* and *STRADA*, collectively delineating a non-proliferative epigenetic landscape characteristic of M-CLL.

Crucially, the genes comprising these three programs were not among the top-ranked features of MOFA, DIABLO, SNF, or NEMO in this cohort. Their identification by PRAMIGO reflects the unique capacity of the heterogeneous graph model to detect features whose relevance emerges from their cross-modal connectivity rather than from their individual marginal effect size, a signal dimension that linear factorization and network fusion methods are not designed to capture.

#### PTB: multi-omics inflammatory programs with metabolite-gene co-activation

In the PTB cohort, the highest-attention subnetwork identified by PRAMIGO revealed an Epithelial-Neural Axis & Tissue Damage module. This subnetwork comprised seven metabolites, including the purine modulator inosine, the kaurenic acid, and signaling lipids lynestrenol and cannabidiolic acid, which were strongly associated with a subset of PTB samples and co-activated with the γδ T-cell marker *TRGV5P* (**Fig. 5D**). This suggests a specific neuro-immunological response at the site of lung injury.

To characterize broader cross-modal structure, we extracted the top 100 genes and 100 metabolites ranked by attention score and performed Leiden clustering^32^ on the induced subgraph. This analysis identified nine coherent multi-omics programs describing distinct biological processes associated with PTB (**Fig. 5E** and **Supplementary Table S5**). Representative programs included: (i) a humoral immune response module enriched for immunoglobulin genes (IGHV family) and the plasma cell regulator *MZB1* together with complement component *C1QC*; (ii) a metabolic reprogramming program linking glycolytic regulator *PFKFB3* with long-chain acylcarnitines; (iii) an oxidative stress and lipid mediator program containing oxidized fatty acids and membrane lipids; (iv) an inflammatory signaling module co-activating *TNFSF15* with arachidonic acid; and (v) an extracellular matrix remodeling program involving structural genes such as *LAMA2* and *MAP1A*.

But the most relevant one is the Pulmonary Barrier Disruption & Neuromodulation program linking tissue-protective genes (*AREG, TRGV5P* and *HLA-V*) with immunomodulatory metabolites (cannabidiolic acid, lynestrenol, and kaurenic acid) in association with neuro-epithelial regulators (*CNTNAP3C, PRKG2* and *LINGO2*) and a dense cluster of polyunsaturated membrane lipids (**Fig. 5F**). These metabolite-gene associations connected epithelial repair and immune surveillance with phospholipid remodeling processes involved in lung barrier integrity and inflammatory adaptation during PTB. By explicitly modeling feature-level cross-modal interactions, PRAMIGO uncovered a biologically coherent homeostatic network that conventional sample-level integration approaches are unable to resolve.

## Discussion

BiomiX 3.0 addresses a fundamental tension in multi-omics research: the growing availability of advanced integration methods on one hand, and their limited accessibility to researchers with limited programming experience. By incorporating five methodologically diverse integration approaches within a single graphical user interface (GUI) platform, BiomiX 3.0 enables systematic, reproducible multi-omics analysis for a broad scientific community without requiring programming expertise. BiomiX 3.0 represents a substantive expansion of the democratization effort initiated with BiomiX v1^17^.

The comparative analysis presented here reveals a central finding with direct practical implications: no single integration method captures the complete biological landscape of a multi-omics dataset. This conclusion, previously supported only by statistical benchmarking metrics, is now substantiated by systematic biological annotation of shared and method-unique features across both the CLL and PTB cohorts. The five methods evaluated: MOFA, DIABLO, SNF, NEMO, and PRAMIGO, identified partially overlapping but substantially distinct molecular signals, and these differences are biologically interpretable rather than merely statistical artifacts.

The CLL results illustrate how methodological diversity translates into biological complementarity. The convergence of all five methods on *EBF1, DIP2C*, and a panel of CpG probes, several of which are established components of validated IGHV stratification panels is particularly informative: it demonstrates that independent analytical frameworks operating on different statistical principles, converge on the same clinically validated epigenetic markers^22^. This cross-method validation strengthens the biological reliability of these features beyond what any single method could provide. The dominance of epigenetic features in the shared signal reflects the well-established primacy of DNA methylation reprogramming as the molecular basis of IGHV-defined CLL subgroups and confirms that this signal is sufficiently strong to be captured regardless of supervision status or analytical framework^23,24^. In contrast, method-unique features reveal the distinct biological dimensions accessible only within specific analytical frameworks: DIABLO’s supervised factorization captures a *KRAB-ZNF* transcriptional repressor programme and BCR-linked signalling (*PLD1*) as high-weight co-varying features^25^; MOFA’s unsupervised multi-factor decomposition independently resolves immune checkpoint regulation (*CTLA4, LAIR1*) and sphingolipid metabolism (DEGS1) as orthogonal biological axes; and PRAMIGO’s graph connectivity uniquely identifies BCR repertoire diversity (IGLV2-14, *IGHV2-5*) and ncRNA co-regulation (cg17130457, *EIF2C3*) as structurally central cross-modal features invisible to loading-weight-based approaches.

The PTB results present a biological landscape that differs structurally from CLL in an instructive way. Rather than a single dominant epigenetic axis, the shared signal in PTB is predominantly metabolomic, with an extensive panel of phosphatidylcholines, lysophosphatidylcholines, and amino acid catabolites co-identified by SNF and NEMO. This metabolome-centred convergence reflects the well-documented systemic lipid and amino acid dysregulation of active tuberculosis, driven by M. tuberculosis consumption of host fatty acids and disruption of one-carbon metabolism^26^. The interferon/innate immune transcriptomic signal, while present through *ANKRD22* and *GBP5*, reaches lower cross-method convergence than the metabolomic signal, suggesting that in this dataset the metabolome carries a stronger and more consistently detectable disease signature across methodological frameworks^27,28^. Method-unique features reveal complementary biological dimensions inaccessible to the shared analysis. DIABLO’s unique features provide a coherent picture of M. tuberculosis metabolic exploitation of the macrophage: the acylcarnitine (3,8)-Decadienoylcarnitine is a direct marker of disrupted mitochondrial β-oxidation consistent with mycobacterial fatty acid consumption^26^, *SLC7A7* identifies the arginine transport checkpoint that determines whether macrophages produce antimicrobial nitric oxide or immunosuppressive polyamines, and *NRP1* points to a Treg-mediated immunosuppressive component in the granuloma microenvironment^29–31^. MOFA’s unique features add systemic dimensions that a supervised approach would not prioritise: *SERPING1* captures an inflammation-interferon response axis, while *RUFY4* and *VPS9D1* represent a lysosomal trafficking restoration programme reflecting the host response to mycobacterial phagosome maturation arrest^34^. PRAMIGO’s unique features, *TRGV5P, IGHV5-51, IGLV10-54*, and *AREG*, demonstrate its specific sensitivity to lymphocyte receptor diversity and tissue repair signals encoded as structural properties of the cross-modal interaction graph rather than as high-variance individual features.

A particularly instructive comparative observation is the divergence in how the two supervised methods (DIABLO and PRAMIGO) prioritize biological signals. DIABLO, using sparse canonical correlation analysis, identifies co-varying high-weight features across modalities within a single latent component, making it sensitive to features that co-vary strongly between RNA and metabolomics layers within a factor. PRAMIGO, through heterogeneous graph attention, identifies features whose biological relevance emerges from their network connectivity across modalities, making it sensitive to features that are structurally central in the cross-modal interaction graph, even if their individual loading weights would be modest. This explains why PRAMIGO uniquely identifies immunoglobulin repertoire diversity (a low-variance, high-structural-connectivity feature) while DIABLO uniquely identifies acylcarnitine profiles (a high-variance, factor-specific metabolomic signal).

Taken together, these findings argue strongly for the concurrent application of complementary methods rather than the selection of a single ‘best’ approach, a strategy now facilitated by BiomiX 3.0. The practical recommendation emerging from this analysis is: (i) use supervised factorization (DIABLO) when the primary goal is biomarker discovery with co-variation across omics layers; (ii) use unsupervised factorization (MOFA) when the goal is multi-dimensional coverage of independent biological axes; (iii) use network methods (SNF/NEMO) when the goal is patient stratification sensitive to sharp between-sample biological boundaries; and (iv) use graph-based learning (PRAMIGO) when the goal is cross-modal regulatory mechanism discovery, particularly when ncRNA, repertoire diversity, or structural cross-modal features are of interest.

PRAMIGO, the novel Heterogeneous Graph Transformer introduced here, occupies a distinct niche among the five methods. Its supervised architecture achieves strong condition-discrimination metrics comparable to those of DIABLO, confirming its effectiveness as a biomarker discovery tool. However, PRAMIGO’s key differentiating capability lies in its explicit modeling of cross-modal molecular interactions through heterogeneous graph edges. The three epigenetic co-regulation programs identified in M-CLL, centered on B-cell lineage transcription factors, prognostic CpG markers, and a broad epigenetic maturity signature, were largely not among the top-ranked features of the other four methods, with individual exceptions such as *EBF1* appearing in the shared set without the cross-modal co-regulation context resolved by PRAMIGO. Similarly, the metabolite-gene co-activation modules identified in PTB provide a level of cross-modal mechanistic specificity that is inaccessible to methods that summarize omic variation at the sample level. PRAMIGO is not presented as a replacement for existing integration approaches, but as a complementary framework designed to address biological questions involving cross-modal molecular regulation. Its heterogeneous graph architecture enables the discovery of interpretable multi-omics programs and feature-level interactions that remain difficult to resolve using conventional sample-level integration methods.

The implementation of the presented methods in the BiomiX 3.0 framework enables the expansion of reproducible pipelines that do not require command-line expertise, instead relying on interactive visualization, automated factor annotation, and bibliography search. No existing platform combines single-omics analysis with five complementary integration methods in a GUI desktop environment accessible to non-bioinformaticians. The expansion of integration in BiomiX follows the ideal of the BiomiX consortium contributors to make these methods more accessible and usable to the scientific community.

BiomiX 3.0 is not without limitations. The platform currently supports transcriptomics, metabolomics, and methylomics, but does not include other omic layers, such as proteomics, genomics (SNP/CNV), or microbiome data. The PRAMIGO architecture requires a minimum number of samples to ensure stable graph construction and supervised contrastive training, which may limit its applicability to very small cohorts. However, graph-based architectures of this class have demonstrated strong performance and scalability in large-scale single-cell datasets^22^. Additionally, while the feature-centric biological analysis presented here provides interpretable insights without relying on underpowered enrichment statistics, it does not replace rigorous pathway analysis when sufficiently powered datasets are available. Future versions of BiomiX will address these limitations by expanding omics compatibility and implementing adaptive sample size handling in PRAMIGO.

By providing non-expert users with access to five methodologically diverse integration approaches within a single, user-friendly environment, and by introducing PRAMIGO as a novel framework for cross-modal molecular interaction modeling, BiomiX 3.0 enables a more complete, multi-perspective interrogation of multi-omics data than has previously been possible without bioinformatics expertise. The comparative framework established here, supported by systematic biological annotation of shared and method-unique features in two disease contexts, provides a principled basis for method selection and interpretation, and the biological findings in CLL and PTB demonstrate the scientific value of deploying multiple complementary integration strategies on the same dataset.

## Conclusions

BiomiX 3.0 expands the BiomiX platform with four new multi-omics integration methods: SNF, NEMO, DIABLO, and the novel graph-based PRAMIGO, within a GUI, user-friendly environment. Benchmarking across two independent cohorts demonstrates that each method captures distinct and partially overlapping biological signals, confirming that method diversity is a scientific asset rather than a limitation. PRAMIGO uniquely models cross-modal molecular interactions through heterogeneous graphs, identifying epigenetic programs in CLL and metabolite-gene co-activation modules in PTB that are inaccessible to conventional approaches. BiomiX 3.0 makes all five methods accessible to researchers without bioinformatics expertise, advancing the democratization of multi-omics integration in line with FAIR data principles.

## Data Availability

The three reference datasets used in the GNN benchmarking, BLCA, LGG, and RCC, are all publicly accessible through the UCSC Xena platform (https://xenabrowser.net/datapages/). The CLL dataset was downloaded from the PACE repository (http://pace.embl.de/) and the PTB dataset from the European Nucleotide Archive (ENA accession PRJNA971365).

## Code Availability

BiomiX 3.0 is publicly available to install and use (https://github.com/BiomiX-consortium/BiomiX3.0). PRAMIGO is inside BiomiX 3.0 and is publicly available also in: https://github.com/iclemente99/PRAMIGO.

## Authors contributions

I.C.L. developed the PRAMIGO methodology, implemented the graph neural network framework, performed the computational analyses, and drafted the manuscript. J.G. designed and implemented the SNF and NEMO workflows in BiomiX, including network construction and visualization, spectral clustering, cluster-number selection and validation, feature prioritization, and clinical enrichment and survival analyses. M.-A.L. and A.G.-d.-l.-F. developed and validated the stabilized Docker-based deployment of BiomiX. C.I. conceived and supervised the study, coordinated software development, and contributed to manuscript preparation. L.V.-S., O.B., N.F., M.H.-V., D.C., and C.J. contributed to data analysis, biological interpretation, software testing, and scientific discussions. A.F.-O. contributed to the design and implementation of the BiomiX integration framework and supervised methodological development. All authors reviewed, edited, and approved the final manuscript.

## Acknowledgement

We also acknowledge the 3TR and PRECISESADS consortium, which supported this work and guaranteed the data used in this article.

## Supplementary Methods

### PRAMIGO architecture

#### Feature selection

For benchmarking analyzes, feature selection followed the procedure described in HeteroGATomics. In the PTB and CLL datasets, PRAMIGO used significantly differential features obtained from single-omic analyzes as graph inputs. This strategy enriches the heterogeneous graph with biologically relevant variables, facilitating the identification of disease-associated cross-modal interaction programs.

#### Autoencoder (AE)

The Autoencoder (AE) layer unifies representations of omic features and samples, producing 256-dimensional embeddings for both entities. These embeddings provide a well-defined node representation when constructing graphs. We implement three separate autoencoders to generate initial embeddings: (1) Omic 1 Autoencoder: Reduces the Omic 1 sample’s representation of each feature to 256 dimensions. (2) Omic 2 Autoencoder: Reduces the omic 2 to 256 dimensions. (3) Sample Autoencoder: Reduces the omic 1 and 2 concatenated features’ dimensional representation of each sample to 256 dimensions. Thus, every cell and gene is assigned an initial 256-dimensional representation.

The loss function for the autoencoder is defined as the Mean Squared Error (MSE) between *X* and its reconstruction 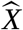:

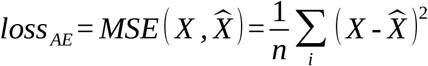

Where *n* is the number of samples. Once we have the embeddings for all features and samples, we can start building the graph.

#### Graph construction

A graph is a mathematical structure represented as *G* =(*V* , *E*), where *V* is the set of nodes (the elements that form the graph) and *E* the set of edges connecting them. In this work, we construct a heterogeneous graph that incorporates multiple node and edge types to capture biological interactions comprehensively .

- Heterogeneous graph: A graph containing multiple types of nodes and/or edges. We define it as *G* =(*V* , *E* , *A* , *R*), where we have the *V* represents nodes, *E* represents edges, *A* represents the node type union, and *R* represents the edge type union.
- Node and edge type mapping functions: We define *τ* (*v*) : *V→ A* and *ϕ* (*e*) : *E→ R* as the mapping functions for node types and edge types, respectively.
- Node meta relation: For a node pair *v*_1_ *v*_2_ linked by an edge *e*_1,2_, the meta relation between *v*_*i*_ and *v* _*j*_ is denoted as ⟨ *τ* (*v*_*i*_) , *ϕ* (*e*_*ij*_) , *τ* (*v* _*j*_) ⟩ .

Our approach constructs an Undirected Binary Heterogeneous Graph with at least three hierarchical levels: samples, omic 1 features, and omic 2 features. Those are the three different types of nodes of the graph. The AE embedding output defines each node .

Within-omic connections are inferred using Pearson correlation, while inter-omic relationships are derived from K-nearest neighbor graphs based on Euclidean distance.

#### Within-omic connections

We used Pearson correlation between the AE embeddings of the same omic features to infer the connections. Pearson threshold is set between 0.7 and 0.9. Everything greater than the threshold is a connection. This way, we infer omic1-omic1 and omic2-omic2 connections.

#### Inter-omic relationships

We used Euclidean distance between the AE embeddings of different node types to infer the connections. The K-nearest neighbor graph connects each embedding to its top K nearest neighbors. K is usually set to about 30% of the distance calculated for each embedding. This way, we infer omic1-omic2, omic1-samples, and omic2-samples connections.

HGT leverages node- and edge-type awareness to transmit information across the heterogeneous graph, while a supervised contrastive loss encourages separation of sample groups in the embedding space.

#### Heterogeneous Graph Transformer

The HGT is employed here in a supervised fashion to learn expressive embeddings for samples and omics, and to estimate their mutual influence via attention mechanisms. Its input is the AE’s latent representation and the constructed heterogeneous graph. Its output is refined embeddings and attention scores that reflect biological relevance.

- Target node and source node: In the context of HGT, a target node *v*_*t*_ is the node currently being updated, while a source node *v*_*s*_ is any neighbor connected to it via an edge *e*_*s,t*_. These roles are crucial for directed attention and message passing.
- Neighborhood graph of target node: For each *v*_*t*_ , a local subgraph *G’* =(*V ‘* , *E ‘* , *A ‘* , *R ‘*) is created, which includes the target, its neighbors *N* (*v*_*t*_), and associated node types *A ‘* and edge types *R ‘*. This local structure allows the transformer to focus on relevant context.

#### Multi-head attention mechanism and vector linear mapping

Each HGT *l*^*th*^layer uses multi-head attention to learn relationships across different embedding subspaces. The embedding of *v*_*t*_ and *v*_*s*_ on the *l*^*th*^layer are denoted as *H*^*l*^ [ *v*_*t*_ ] and *H*^*l*^ [ *v*_*s*_ ]. Each head *h* learns:

- A query vector: 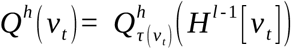
- A key vector: 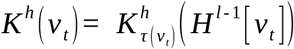
- A value vector: 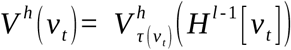

Each node type has its own linear projection layers to adapt to structural heterogeneity.

#### Heterogeneous mutual attention

The mutual attention between *v*_*t*_ and *v*_*s*_ is calculated with a specialized attention operator:

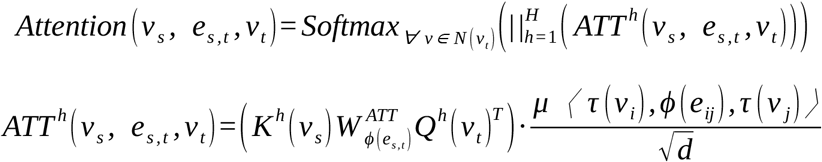

Where *d* is the dimension of the embedding and *μ* is a learnable scalar denoting the importance of the meta-relation (node-type, edge-type, node-type).

#### Heterogeneous message passing

Each source node passes a message to the target node:

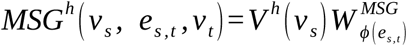

Where 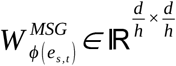 is a learnable matrix for incorporating edge dependency. These are concatenated across all heads:

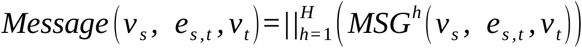

#### Target specific aggregation

The new embedding of *v*_*t*_ at *l*^*th*^ layer is obtained by aggregating all messages weighted by attention:

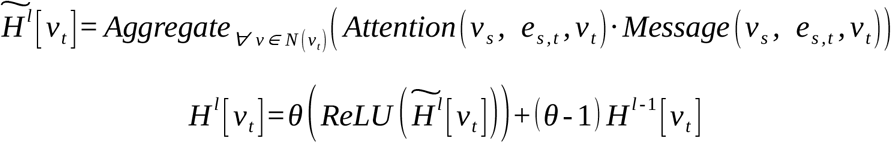

Where *ReLU* is the activation function and θ is a trainable parameter controlling residual blending.

#### HGT training on subgraph

Due to the massive size of the input graph, we apply a subgraph training strategy (HGSampling), selecting a subset of patients per batch based on expression-based sampling:

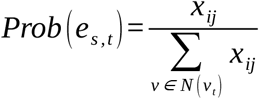

This way *Prob* (*e*_*s* , *t*_) acts as the probability of selecting and edges to be part of the subgraph. Each batch feeds one subgraph to the HGT, and model parameters are sequentially updated.

The model is trained in a supervised representation-learning setting. Instead of reconstructing expression matrices with KL divergence, the training objective combines classification supervision and metric learning on sample embeddings. Specifically, the loss function is defined as a weighted sum of cross-entropy loss and cosine embedding loss:

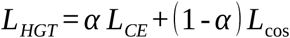

where *L*_*CE*_ is the cross-entropy loss between predicted sample class logits and ground-truth labels, encouraging discriminative sample representations, *L*_cos_ and is the cosine embedding loss applied to pairs of sample embeddings. Positive pairs correspond to samples from the same class. In contrast, negative pairs correspond to samples from different classes, enforcing intra-class similarity and inter-class separation in the learned embedding space.

Training stops either upon convergence, defined as no improvement in the loss over a predefined number of epochs, or after reaching a fixed maximum number of epochs.

### Training losses definitions

#### Cross-entropy loss

*L*_CE_. Let *N* be the number of samples in a mini-batch, *C* the number of classes, *z*_*i*_ ∈ ℝ^*C*^ the predicted logits for sample *i*, and *y*_*i*_ ∈ {1 ,*…, C* } its ground-truth label. The softmax probability of class *c* is

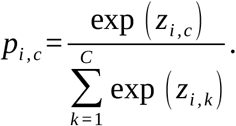

The cross-entropy loss is then

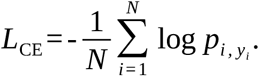

This term encourages the sample embeddings to be linearly separable with respect to the class labels.

#### Cosine embedding loss

*L*_COS_. Let *e*_*i*_ , *e*_*j*_ ∈ ℝ^*d*^ be the embeddings of two samples and let *y*_*ij*_ ∈ {+ , 1 - 1} indicate whether the pair is positive (*y*_*ij*_ =+ 1, same class) or negative (*y*_*ij*_ =-1, different classes). The cosine similarity is

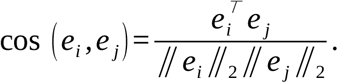

The cosine embedding loss for a pair is defined as

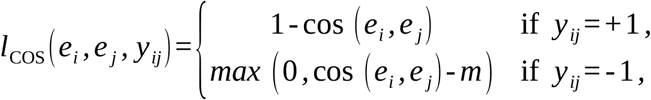

where *m* ∈ [0,1) is a margin hyper-parameter (commonly set to 0). Averaging over all pairs (*i* , *j*) in the batch yields

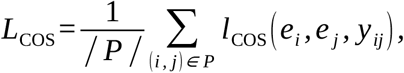

with *P*the set of pairs considered in the batch. Positive pairs are pulled together in the embedding space while negative pairs are pushed apart beyond the margin, thereby enforcing intra-class compactness and inter-class separation.

### Relative omic contribution calculation

The relative contribution of each omic layer is method-specific and derived directly from the native weighting/importance quantities produced by each algorithm; these are then normalized (where needed) to allow comparison of relative omic importance.

#### DIABLO

DIABLO (block.splsda / sGCCA framework in mixOmics) constructs latent components that maximize the sum of covariances between blocks (weighted by the design matrix) while discriminating the phenotype. After model fitting, the proportion of variance explained by each component in each omic block is obtained from the model object (prop_expl_var).

For a given component *c*and omic block *q*, this is the fraction of total variance in the (centered/scaled) block *X*^(*q*)^captured by the component scores 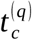 :

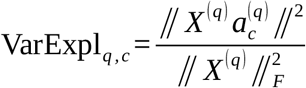

(where 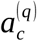 are the loadings). The global omic contribution is obtained by summing (or averaging) VarExpl_*q* ,*c*_across the retained components. This quantifies how much of the variation in each omic is captured by the shared discriminative components.

#### MOFA

MOFA decomposes each omic view *m*as *Y* _*m*_ *≈W*_*m*_ *Z* + noise. After training, the variance explained by factor *k*in view *m*is the coefficient of determination:

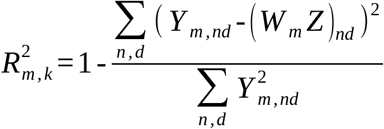

(or the equivalent Gaussian pseudo-data version for non-Gaussian likelihoods).

The total variance explained by omic *m*is 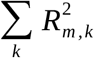 (or the joint *R*^*2*^ across all factors). The relative contribution of each omic is therefore the fraction of total reconstructed variance attributable to that view across the active factors. This is exactly the quantity returned by get_variance_explained() / calculateVarianceExplained().

#### SNF and NEMO

Neither method produces a direct “variance explained” quantity. Instead, feature-level importance is computed via Normalized Mutual Information (NMI) with the fused similarity network (or the resulting clustering), following the approach implemented in SNFtool::rankFeaturesByNMI.

For each feature *f* belonging to omic *q*:

1. Discretize/cluster the samples according to the values of *f* .
2. Compute the Normalized Mutual Information between this clustering and the consensus network *W* obtained by SNF (or the equivalent neighborhood-based consensus graph of NEMO):

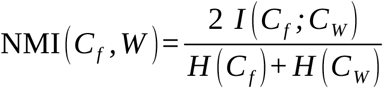

where *I* is mutual information and *H* is entropy.

The top 100 features per omic (ranked by NMI) are retained. The omic contribution is then the sum (or mean) of the NMI scores of those top-100 features belonging to omic *q*, optionally normalized by the total NMI mass across all omics so that the contributions sum to 1. This measures how strongly the features of each omic agree with the multi-omics consensus structure.

#### HGT (Heterogeneous Graph Transformer)

HGT assigns multi-head attention scores to edges/nodes in the heterogeneous multi-omics graph. For each omic feature node *v*the attention coefficient *α*_*u,v*_(or the aggregated attention mass arriving at / leaving from *v*) quantifies its importance in the message-passing that produces the final sample embeddings.

The omic contribution is the sum of the absolute attention scores of all feature nodes belonging to that omic (or the average attention per feature of that omic), again optionally normalized across omics. This directly reflects the relative influence each omic layer exerts on the learned representations.

#### Feature-overlap analysis

Independently of the omic-level contribution, feature-level concordance is assessed with the Jaccard index on the top-20 features of each omic (ranked by absolute loading weight for DIABLO/MOFA, by NMI for SNF/NEMO, or by attention score for HGT):

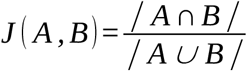

where *A*and *B*are the sets of top-20 features returned by two methods for the same omic layer. The choice of 20 features is a fixed hyper-parameter in BiomiX 3.0 chosen for interpretability.

### PARMIGO’s hyper-parametrization

For PRAMIGO, graph construction and training of the Heterogeneous Graph Transformer (HGT) model, the architecture was configured with a hidden dimension of 128, an autoencoder hidden dimension of 256, 8 attention heads, 2 GNN layers, and a dropout ratio of 0.0 using the ‘hgt’ layer type and an embedding node autoencoder (AEtype 1). Graph construction utilized Pearson correlation cutoffs of 0.7 for both omic feature sets, a K-nearest neighbors metric set to *k* = round (samples.shape [0 ] / 3) using cosine similarity, a Leiden clustering resolution of 1.5, and a minimum cluster size of 3 features. Training was performed over 100 epochs with 25 sub-sampled graphs per epoch, a batch size of 32, a learning rate of 0.0001 using the AdamW optimizer, a regularization weight of 0.0, and a supervised loss function. A ReduceLROnPlateau learning rate scheduler was integrated with an attenuation factor of 0.5 and a patience of 5, with checkpoints saved every epoch. Data preprocessing maintained a fixed random seed of 0 for reproducibility, set omic sampling rates to 1.0, and partitioned the data into 70% training, 15% validation, and 15% test splits.

## Supplementary Figures

**Supplementary Fig. 1:**
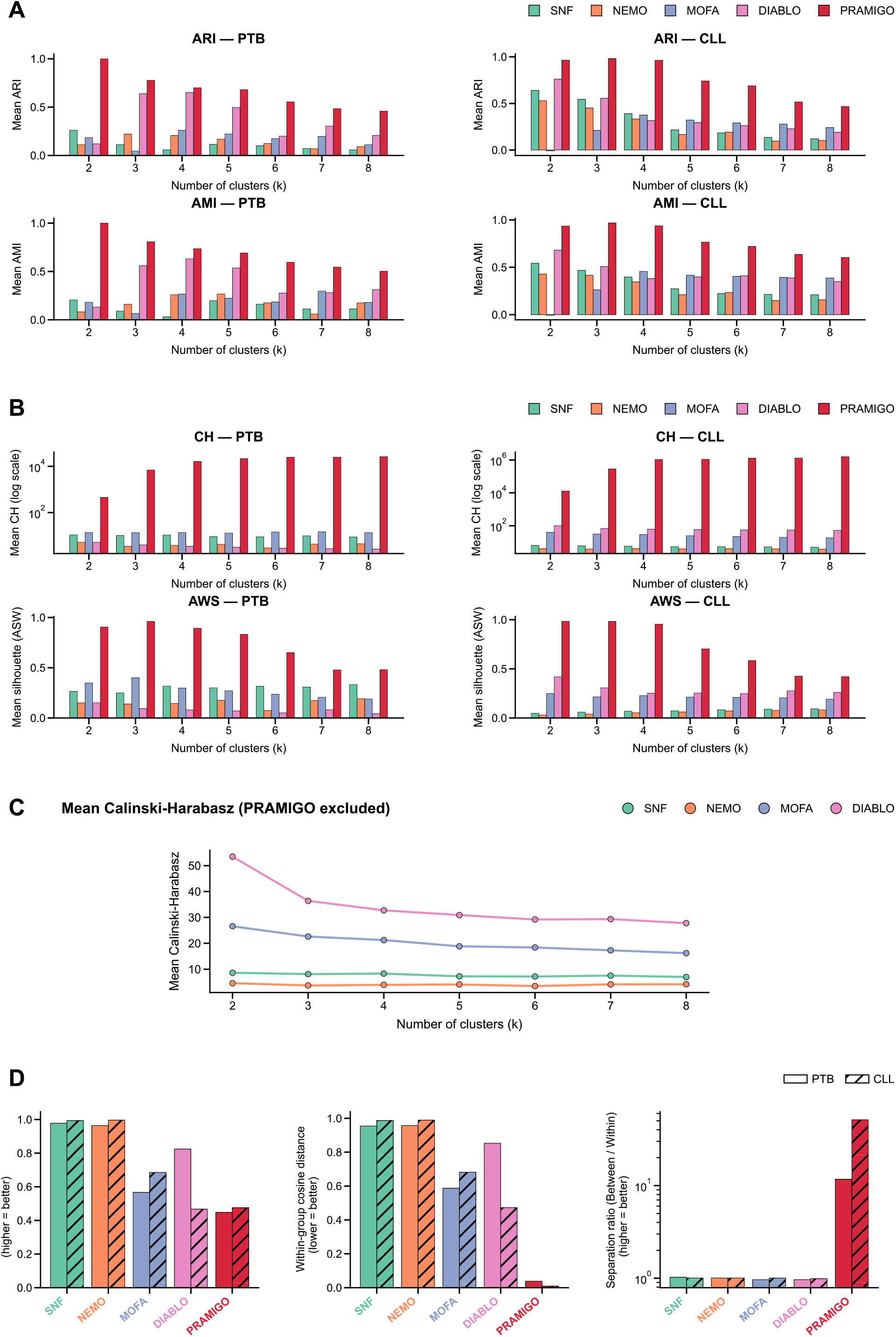
Detail understanding of PRAMIGO programs. (A) Extended Adjusted Rand Index (ARI) and Adjusted Mutual Information (AMI) for all five methods on both cohorts. (B) Extended Average silhouette width (ASW) and Calinski-Harabasz (CH) index for all five methods on both cohorts. (C) Calinski-Harabasz (CH) index for all five methods on both cohorts excluding PRAMIGO (D) Inter/intra-condition distance ratio in latent space.

**Supplementary Fig. 2:**
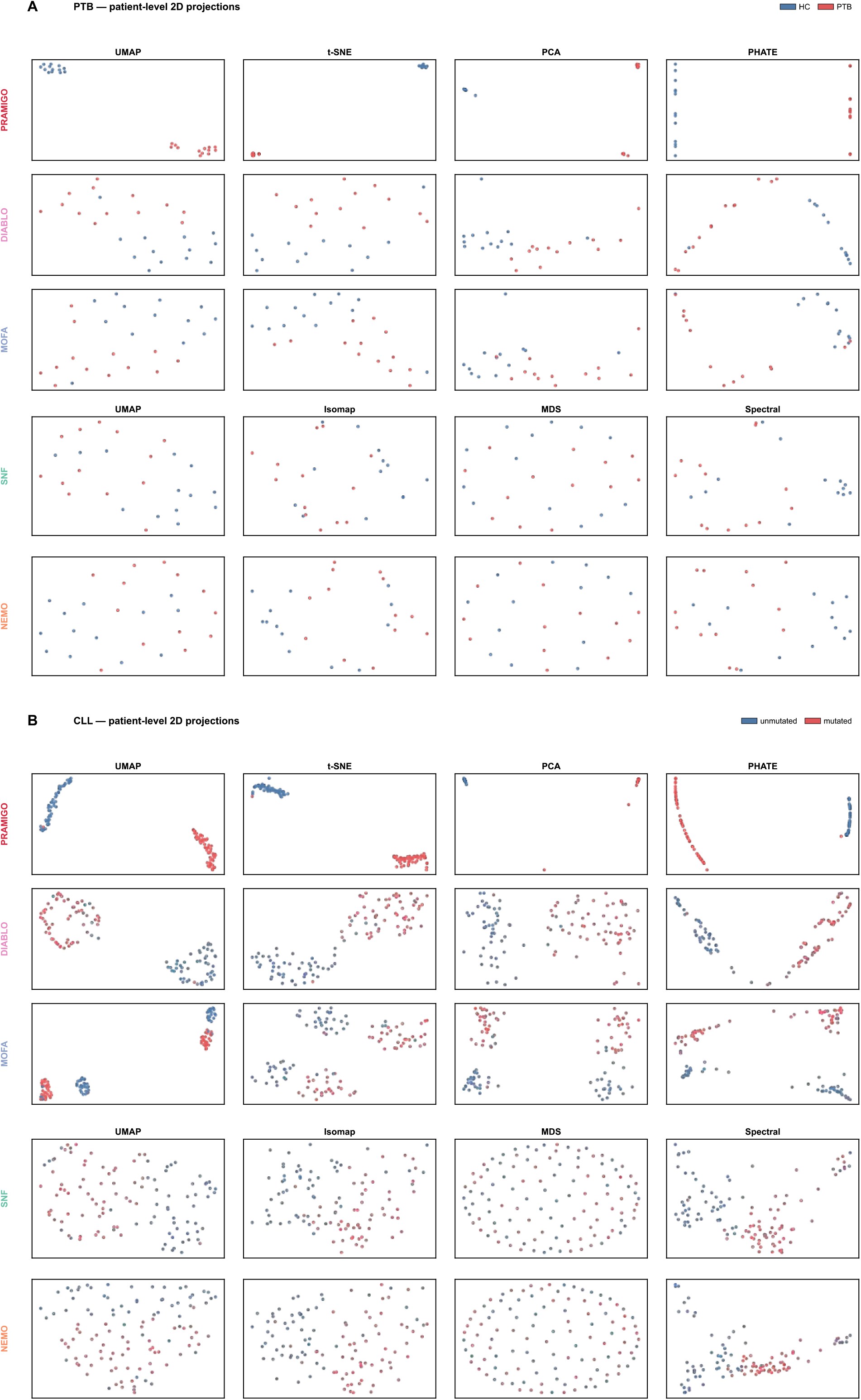
Complete visualization of BiomiX 3.0 methods. (A) Full visualization panel of PTB samples with PRAMIGO, DIABLO and MOFA with UMAP, PCA, tSNE, and PHATE, while NEMO and SNF with UMAP, MDS, Isomap and Spectral colored by condition. (B) Full visualization panel of CLL samples with PRAMIGO, DIABLO and MOFA with UMAP, PCA, tSNE and PHATE, while NEMO and SNF with UMAP, MDS, Isomap and Spectral colored by condition.

**Supplementary Fig. 3:**
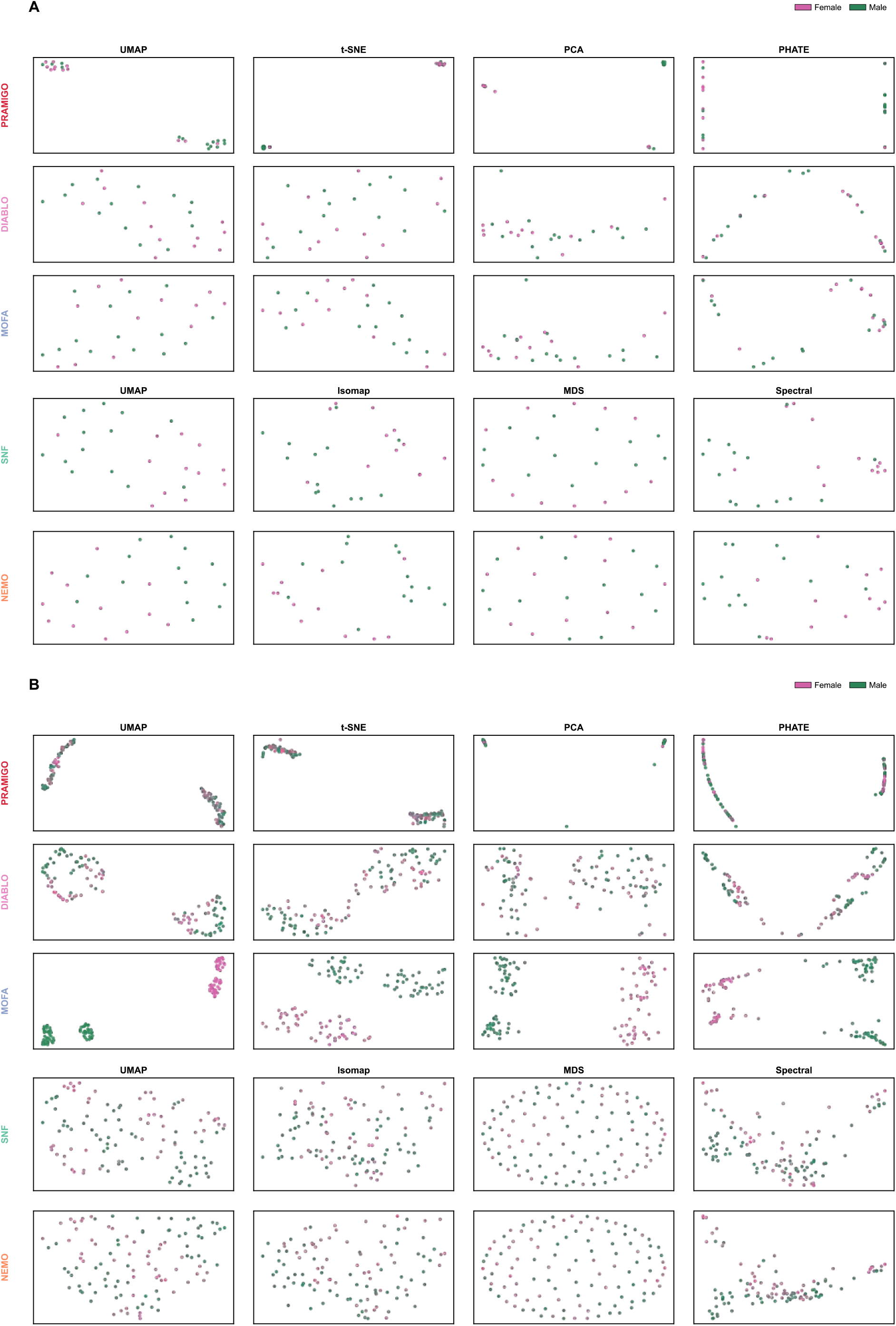
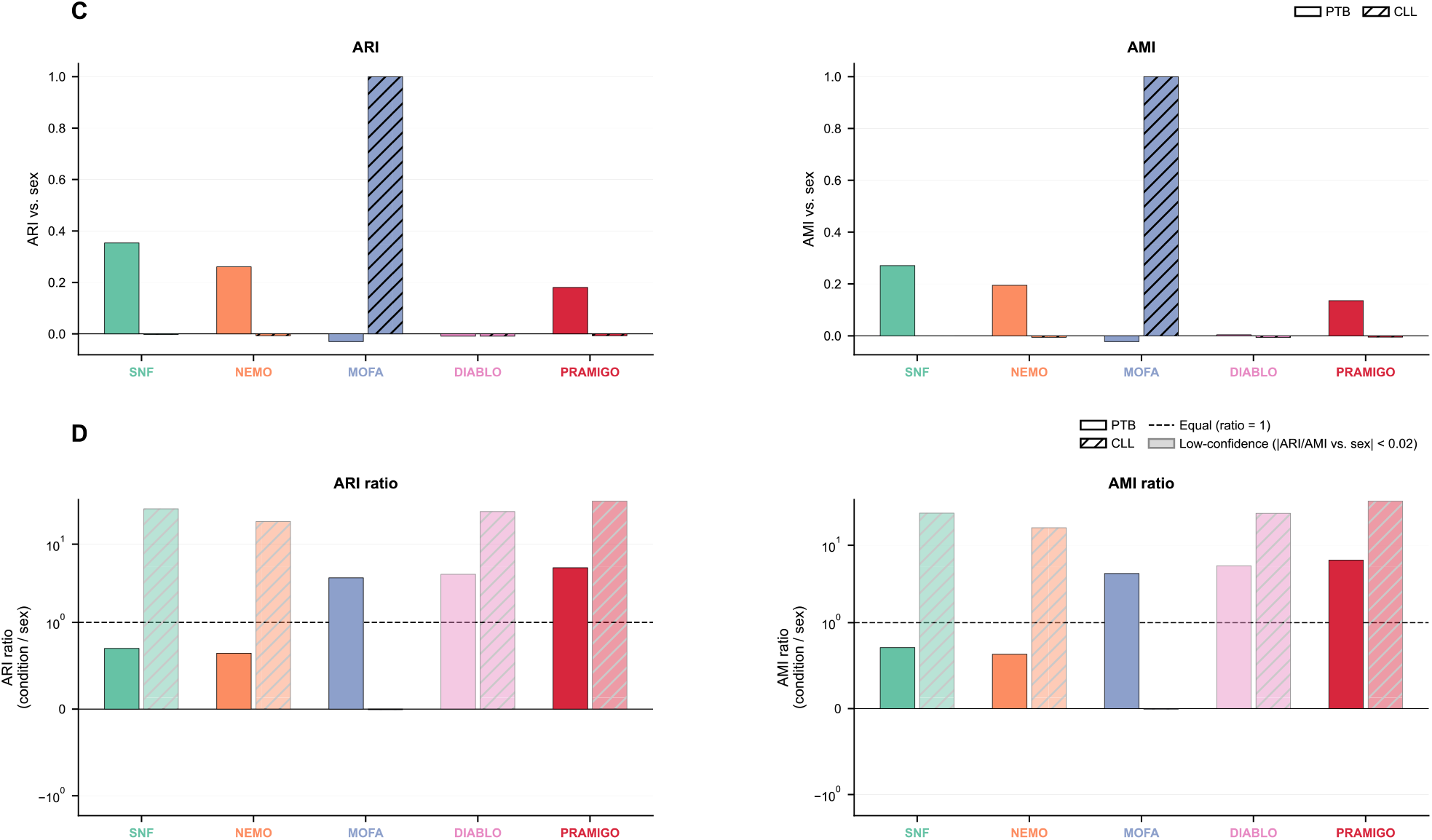
Unsupervised multi-omics representations are driven by sex. (A) Comprehensive visualization of PTB sample representations generated by PRAMIGO, DIABLO and MOFA using UMAP, PCA, t-SNE and PHATE, and by NEMO and SNF using UMAP, MDS, Isomap and spectral embedding. Samples are colored according to sex. (B) Comprehensive visualization of CLL sample representations generated by PRAMIGO, DIABLO and MOFA using UMAP, PCA, t-SNE and PHATE, and by NEMO and SNF using UMAP, MDS, Isomap and spectral embedding. Samples are colored according to sex. (C) Adjusted Rand Index (ARI) and Adjusted Mutual Information (AMI) scores obtained for each method using k = 2 clusters and compared with sex labels. (D) Ratio between the ARI and AMI scores obtained for the condition labels and those obtained for sex labels, quantifying the relative extent to which the resulting sample representations are associated with disease condition versus sex.

**Supplementary Fig. 4:**
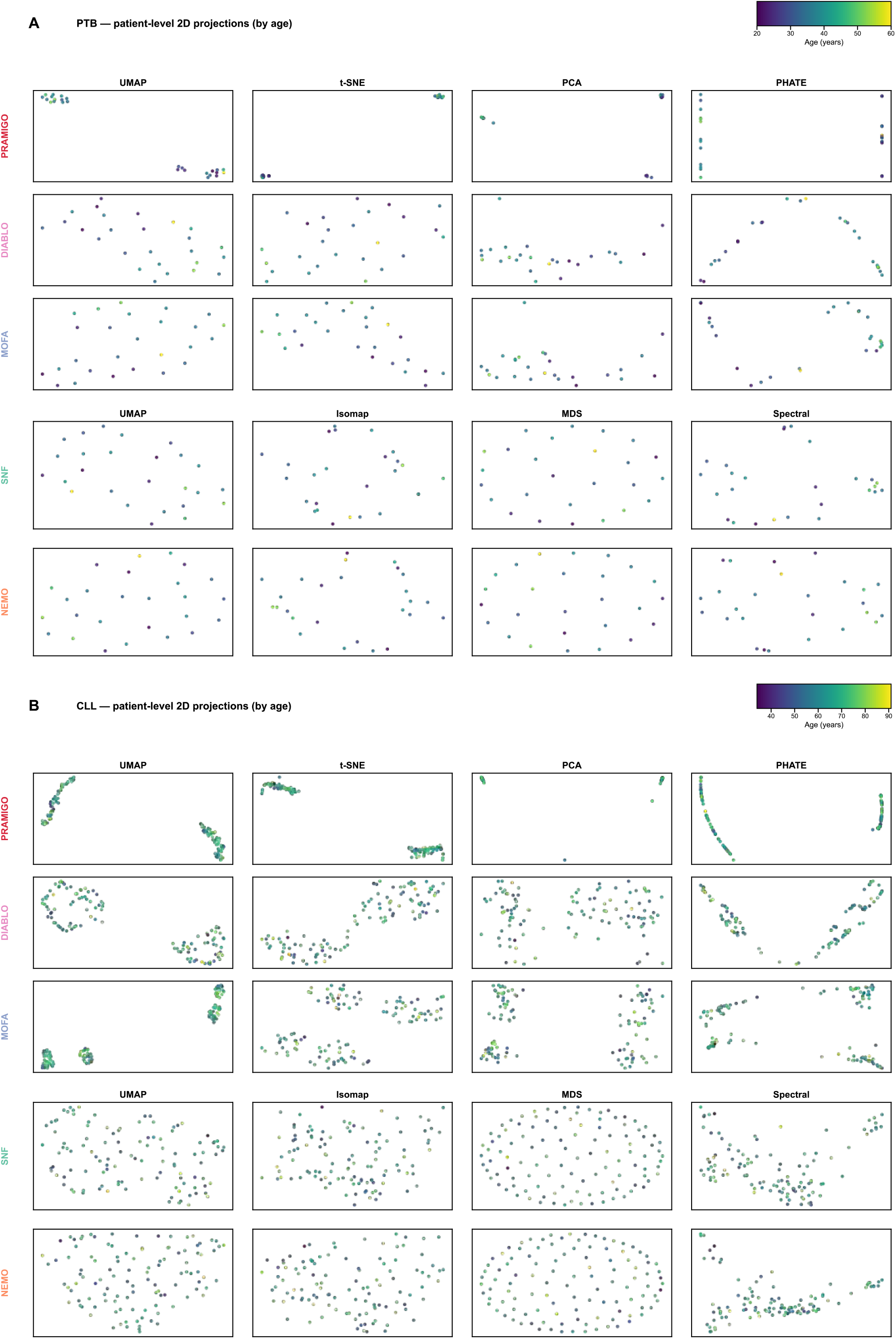
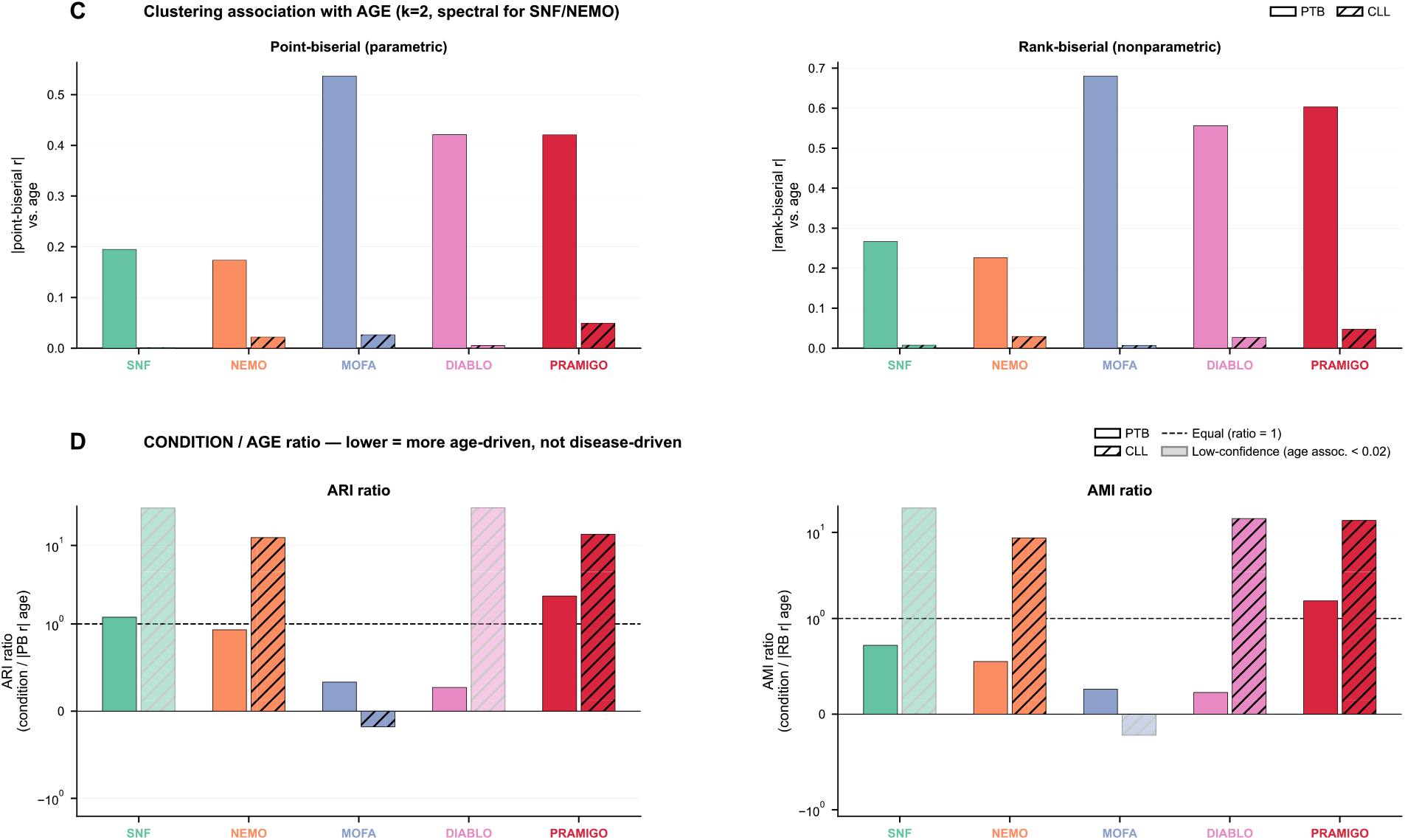
Unsupervised multi-omics representations are driven by age. (A) Comprehensive visualization of PTB sample representations generated by PRAMIGO, DIABLO and MOFA using UMAP, PCA, t-SNE and PHATE, and by NEMO and SNF using UMAP, MDS, Isomap and spectral embedding. Samples are colored according to age. (B) Comprehensive visualization of CLL sample representations generated by PRAMIGO, DIABLO and MOFA using UMAP, PCA, t-SNE and PHATE, and by NEMO and SNF using UMAP, MDS, Isomap and spectral embedding. Samples are colored according to age. (C) Association between each method’s sample representation and age, obtained using k = 2 clusters and quantified with the absolute point-biserial correlation and absolute rank-biserial correlation (derived from a Mann-Whitney U test) between cluster assignment and age; ARI and AMI could not be used directly for this comparison as they require a categorical, rather than continuous, ground truth. (D) Ratio between the ARI and AMI scores obtained for the condition labels and the corresponding point-biserial and rank-biserial correlation scores obtained for age, using the same k = 2 clustering, quantifying the relative extent to which the resulting sample representations are associated with disease condition versus age.

**Supplementary Fig. 5:**
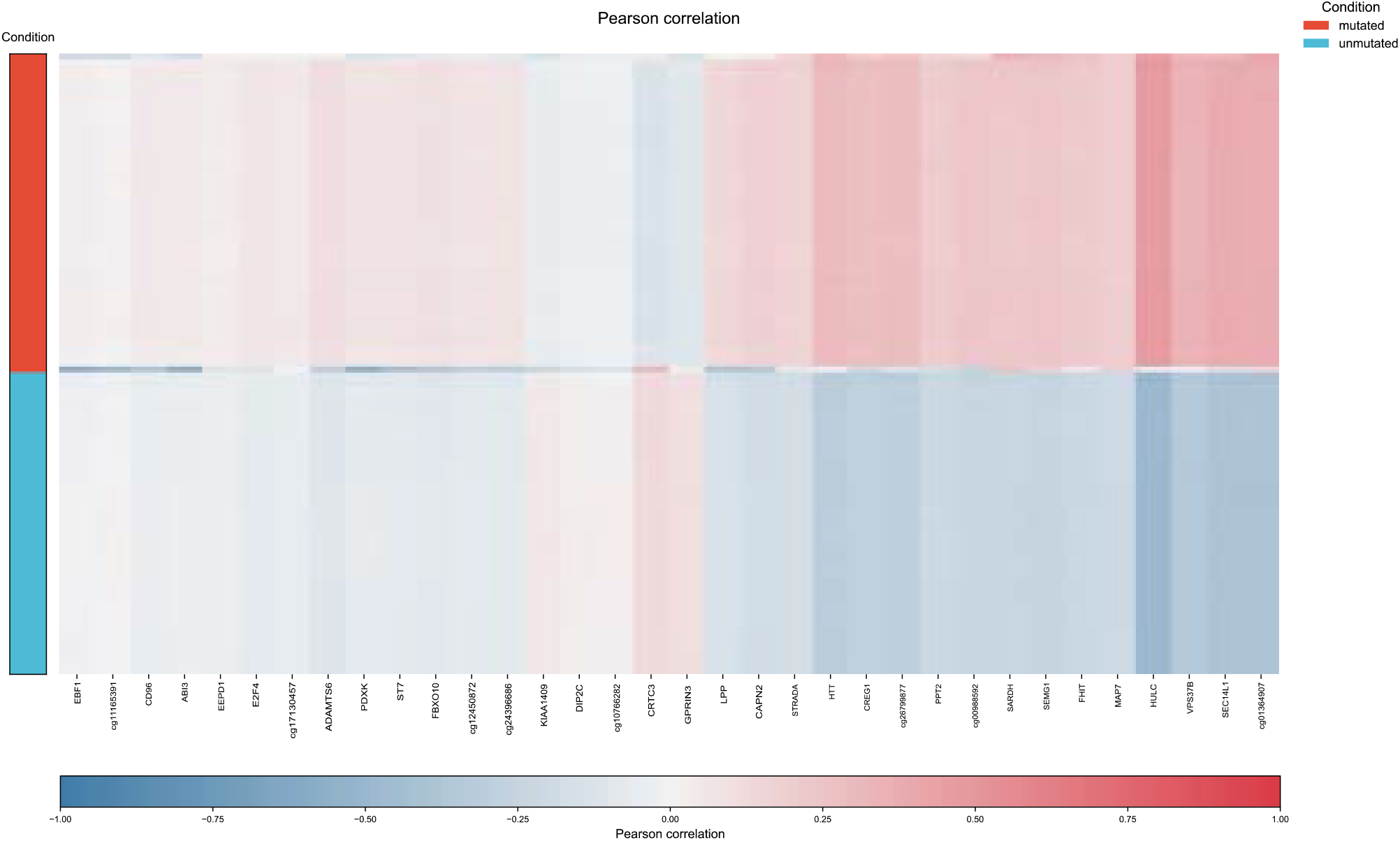
Detail understanding of PRAMIGO programs. Correlation heatmap of the top 34 methylomic genes embeddings with CLL mutated and unmutated sample embeddings.

